# *De novo* human brain enhancers created by single nucleotide mutations

**DOI:** 10.1101/2021.07.04.451055

**Authors:** Shan Li, Sridhar Hannenhalli, Ivan Ovcharenko

**Affiliations:** Computational Biology Branch, National Center for Biotechnology Information, National Library of Medicine, National Institutes of Health, Bethesda, MD, 20892, USA; Cancer Data Science Laboratory, Center for Cancer Research, National Cancer Institute, National Institutes of Health, Bethesda, MD, 20892, USA

## Abstract

Advanced human cognition is attributed to increased neocortex size and complexity, but the underlying gene regulatory mechanisms are unknown. Using deep learning model of embryonic neocortical enhancers, and human and macaque embryonic neocortex H3K27ac data, we identified ~4000 enhancers gained *de novo* in the human, largely attributable to single-nucleotide essential mutations. The genes near *de novo* gained enhancers exhibit increased expression in human embryonic neocortex relative to macaque, are involved in critical neural developmental processes, and are expressed specifically in the progenitor cells and interneurons. The gained enhancers, especially the essential mutations, are associated with central nervous system disorders/traits. Integrative computational analyses suggest that the essential mutations establish enhancer activities through affecting binding of key transcription factors of embryonic neocortex. Overall, our results suggest that non-coding mutations may have led to *de novo* enhancer gains in the embryonic human neocortex, that orchestrate the expression of genes involved in critical developmental processes associated with human cognition.

## Introduction

The neocortex is a mammalian innovation enabling complex cognitive and motor tasks (Geschwind and Rakic 2013; Emera et al. 2016). The substantial expansion and functional elaboration of the neocortex provides an essential basis for the advanced cognitive abilities of humans (Geschwind and Rakic 2013), which includes an increase in the proliferative capacity of the progenitor cells (Dehay et al. 2015; Namba and Huttner 2017; Sousa et al. 2017), an increase in the duration of their proliferative, neurogenic and gliogenic phases (Lewitus et al. 2014; Otani et al. 2016), an increase in the number and diversity of progenitors, modification of neuronal migration, and establishment of new connections among functional areas (Geschwind and Rakic 2013).

Critical events in corticogenesis, including specification of cortical areas and differentiation of cortical layers require precise spatiotemporal orchestration of gene expression (Rakic et al. 2009). Modifications in gene regulation are thus hypothesized to be a major source of evolutionary innovation during cortical development (Rakic 2009; Rakic et al. 2009; Geschwind and Rakic 2013). Among these are gain and loss of enhancers, repurposing of existing enhancers, rewiring of enhancer-gene interaction networks, and modifications of crosstalk between enhancers operating within the same cis-regulatory landscape (Long et al. 2016). However, several fundamental questions remain open: to what extent the evolutionary gain and loss of enhancers has contributed to human-specific features of corticogenesis? Specifically, how often enhancer gain is associated with an increased expression of the target gene involved in human corticogenesis? To what extent the emergence of human-specific enhancers could be explained by a single or a few single-nucleotide mutations? How often do such mutations establish an enhancer from neutral DNA through creation of binding sites of activators as opposed to the disruption of binding sites of repressors? What are the transcription factors (TFs) mediating critical enhancer gains and losses and what gene regulatory networks are induced by such mutations? A previous study identified Human Gained Enhancers (termed HGEs) (Reilly et al. 2015) that exhibit increased regulatory activity in human relative to macaque and mouse. In contrast, our focus is *de novo* gained enhancers in human that presumably originate from neutral non-coding sequence via minimum number of single-nucleotide substitutions along the human lineage. Besides the availability of enhancer activity profiles in the developing brain of humans and macaques (Reilly et al. 2015), a quantitative model that can accurately estimate enhancer activity from DNA sequence, with single-nucleotide sensitivity, is critical to answering the questions above.

In this study, we developed a deep learning model (DLM) able to learn the sequence encryption of human and primate embryonic neocortex enhancers, enabling us to quantify the functional effect of single-nucleotide mutations on enhancer activity. Leveraging the DLM and the recently available enhancer activity profiles in developing neocortex in humans and macaques (Reilly et al. 2015), we identified single-nucleotide mutations that potentially drive human-specific regulatory innovations. We observed that a single-nucleotide mutation is often sufficient to give rise to an enhancer, leading to increased expression of the proximal target gene. As a group, *de novo* gained enhancers induce genes that are critical to cognitive function and are expressed preferentially in the progenitor and interneuron cells of the developing neocortex. *De novo* gained enhancers and their target genes induce and mediate a potential core regulatory network in the developing human neocortex, with POU3F2 occupying a central position. Essential single-nucleotide mutations resulting in *de novo* enhancer gain exhibit relaxed negative, or potentially adaptive, selection. Interestingly, the essential mutations and *de novo* gained enhancers are enriched for cognitive traits; in particular, the *de novo* gained enhancers associated with regulation of key TFs are enriched for *de novo* mutations in patients with the autism spectrum disorder (ASD). Compared to HGEs, although *de novo* gained enhancers have relatively weaker enhancer activity, they are more likely to turn on gene expression in human and regulate genes associated with brain development. Integrating a DLM with epigenomic data allowed us not only to identify *de novo* gained human-specific enhancers that might underlie advanced cognition, but also gauge the impact of single-nucleotide mutations in this process.

Overall, our results, based on the H3K27ac profiles in developing human and macaque brain, and a novel sequence-specific deep learning model of embryonic neocortical enhancers, suggests a wide-spread *de novo* gain in enhancers, largely driven by single nucleotide mutations, in the progenitors and interneurons of the developing human neocortex, that together induce a core regulatory network that associated with human cognitive abilities as well as cognitive disorders.

## Results

### Identifying de novo enhancer gain and essential human mutations - Overview

To assess functional impact of single nucleotide mutations on enhancer activity, we leveraged the H3K27ac ChIP-seq data during human and macaque corticogenesis as a proxy for active enhancers (Reilly et al. 2015) and built a DLM to learn the regulatory code encrypted in the enhancer sequences (Figure S1A-C, Methods). Next, by integrating the predicted enhancer activities in human, macaque, and the human-macaque common ancestor inferred from multiple sequence alignment (Paten et al. 2008) based on a probabilistic model (Holmes and Bruno 2001; Holmes 2003; Bradley and Holmes 2007) with the observed enhancer activities in human and macaque, we identified human-specific *de novo* gains and losses of enhancers (Figure 1A, Methods). We then prioritized the single-nucleotide human-macaque mutations in the *de novo* gained and lost enhancers based on the difference of the DLM scores between the macaque sequence and the intermediate sequence with one or more introduced human allele(s). For an enhancer with multiple mutations, which was either gained or lost in the human genome, we first introduced each human-specific allele to its matching macaque sequence and estimated its impact on enhancer activity using the difference in the DLM score attributed to the human allele. By iteratively increasing the number of introduced human-specific alleles and scoring the modified sequence, we evaluated the impact of combinations of mutations and determined the minimal number of mutations needed for an enhancer to be gained or lost in the human lineage.

**Figure 1.**
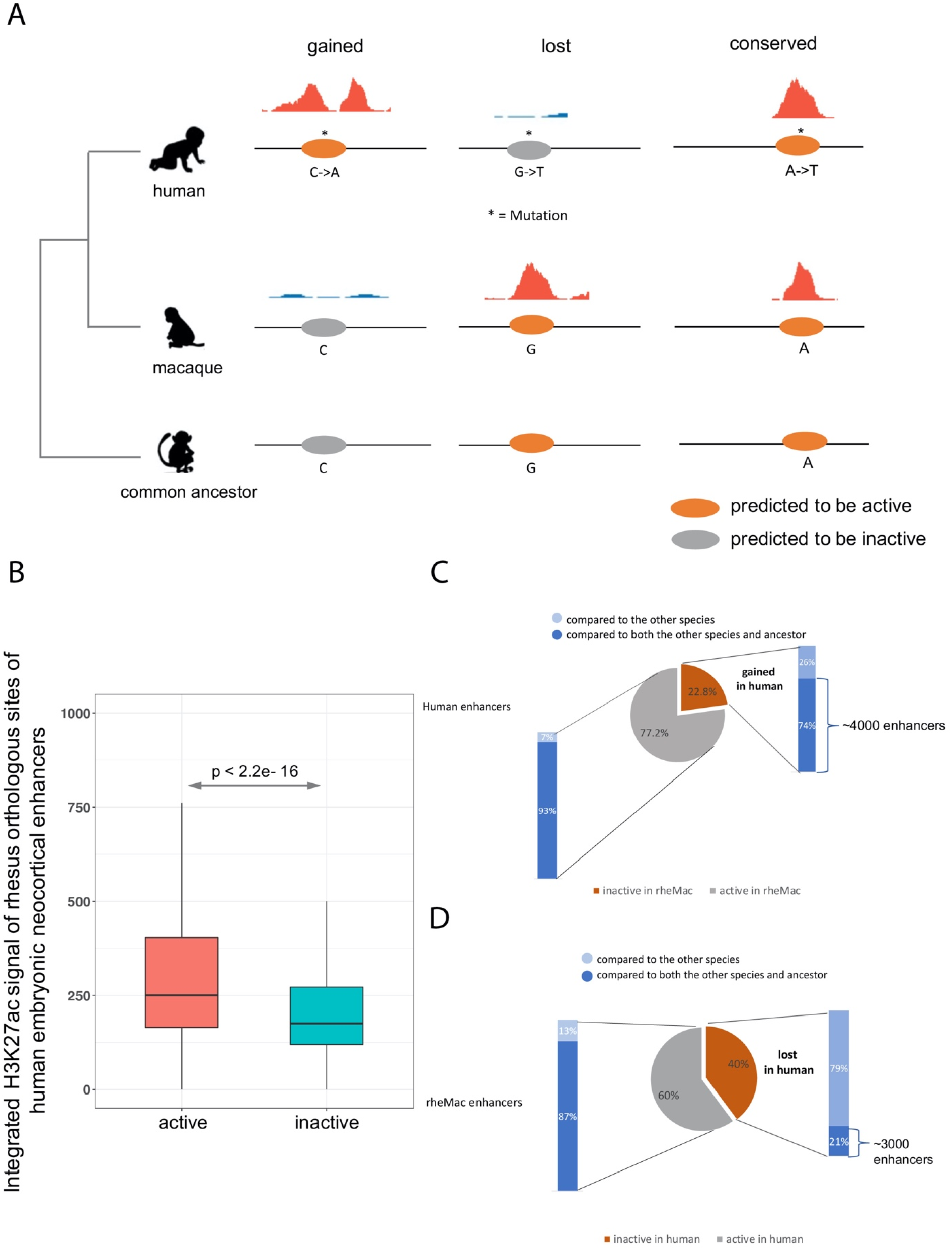
*De novo* gained and lost enhancers. A) Identification of *de novo*-gained, lost, and conserved enhancers. If a human enhancer scored highly by the DLM and scored low both in macaque and in the common ancestor, and was not detected by H3K27ac in macaque, it was considered to be gained in humans. If a macaque enhancer having high DLM score, scored high in common ancestor, scored low in human and was undetectable by H3K27ac in human it was considered a loss in human. The enhancers that are detected by H3K27ac in both human and macaque and scored highly in all three genomes were called conserved enhancers. B) Comparison of embryonic macaque neocortex integrated H3K27ac signal intensities (within the 1kb enhancers) between the predicted active and inactive macaque orthologs of human embryonic neocortex enhancers. C) The fraction of *de novo* gained human embryonic neocortex enhancers by comparing human to both macaque and their common ancestor. Specifically, 74% of human enhancers that are inactive in macaque are active in the common ancestor and 93% of human enhancers that are active in macaque are active in the common ancestor. D) The fraction of lost human embryonic neocortex enhancers by comparing human to both rhesus macaque and their common ancestor. Specifically, 21% of macaque enhancers that are inactive in human are active in the common ancestor and 87% of macaque enhancers that are active in human are active in the common ancestor. Light blue refers to relative to the other species, dark blue refers to relative to both the other species and common ancestor.

### An accurate DLM of embryonic neocortex enhancers in human and macaque

The human embryonic neocortex H3K27ac ChIP-seq peaks were obtained from the four temporal/spatial groups: the whole cortex at 7 post conception weeks (p.c.w.) (CS16) and 8.5 p.c.w. (CS23) and primitive frontal and occipital tissues from 12 p.c.w. (F2F and F2O) (Reilly et al. 2015). We trained a DLM separately for each set of enhancers (Methods). The DLM was able to discriminate human embryonic neocortex enhancers from accessible regions devoid of non-fetal-brain-enhancer with high accuracy: the area under the receiver operating characteristic curve (auROC) ranges from 0.9 to 0.94 (Figure S1B), and the area under the precision-recall curve (auPRC, expectation = 0.091) ranges from 0.56 to 0.63 for the four datasets (Figure S1C). The consistently high accuracy of all models showed the ability of DLMs in capturing sequence signatures of brain enhancers similarly to previous modeling of enhancers in other cells and tissues (Supplementary Results 1), and prompted us to conjecture that the four groups of enhancers tend to share either genomic locations or sequence characteristics. To assess their sequence similarity, we trained the DLM on one set and predicted those from all other sets. We observed both high auROCs and auPRCs (Figure S1D), strongly suggesting a shared sequence characteristics across the four enhancer sets. However, the genomic overlap between any two groups of enhancers is relatively low (20-40%) (Figure S1E), indicating that the four sets of enhancers overlap only partially but share sequence characteristics.

We proceeded to investigate the *de novo*-gain and loss of enhancers by comparing human 8.5 p.c.w (CS23) sample and macaque sample at approximately matching time point (e55) (Reilly et al. 2015), as the DLM trained on CS23 has not only high auROC (0.92) but also the highest precision at a low false positive rate (FPR = 0.1) (Figure S1BC). To ascertain that the DLM trained on CS23 can accurately predict the enhancer activity in macaque, we scored the macaque orthologs of CS23 enhancers and compared the e55 H3K27ac signal intensities of the macaque orthologs predicted to be active with those predicted to be inactive (Methods). The predicted active regions indeed have significantly stronger H3K27ac signal (Figure 1B), suggesting that the DLM learned from human embryonic neocortical enhancers can accurately gauge the enhancer activity in macaque from its genomic sequence.

We next identified the enhancers *de novo*-gained, lost, or conserved in human relative to both macaque and human-macaque common ancestor based on the H3K27ac profile and DLM scores (Methods, Figure 1A). In total, we identified 4,066 *de novo* gained (Figure 1C), 2,925 lost, and 23,119 conserved neocortical enhancers (Figure 1D). Although the majority of the developmental neocortical enhancers remained active since the divergence of human and macaque from their common ancestor, there are certain groups of enhancers that are gained or lost in the human lineage, prompting us to conjecture that these gain and loss events may correlate with the human-specific features of corticogenesis, which we investigate next.

### *De novo* gained enhancers are associated with critical cortical developmental functions

Next, to investigate whether *de novo* enhancer gains are accompanied by an increase in the expression of their putative target genes, we compared the human-to-macaque ratios of gene expression near gained enhancers versus those near lost enhancers and observed that the genes near gained enhancers show a human-specific increase in expression while a reverse trend is exhibited by genes near lost enhancers (Figure 2A); this trend holds when we rely on Hi-C contact data to map an enhancer to its target genes (Figure S2). Consistently, gained enhancers are enriched near the genes with top 5% highest expression relative to macaque (Figure 2B). Notably, the fetal brain eQTLs (O’Brien et al. 2018) are significantly enriched in *de novo* gained enhancers compared to lost and conserved enhancers (Figure 2C and Figure S3). These results together support a causal link between enhancer gain and an increase in the expression of their target genes. Furthermore, the *de novo* gained enhancers are primarily associated with gliogenesis, neural tube development, and neural precursor cell proliferation, among other central nervous system (CNS) related developmental processes (Figure 2D, Figure S4A, and Table S1). In contrast, lost enhancers are associated with only a small number of CNS related essential biological processes, including regulation of axon extension, neural retina development, neural precursor cell proliferation, and cerebral cortex cell migration (Figure 2E, Figure S4B, and Table S2). Lost enhancers are enriched for far fewer processes than the *de novo* gained enhancers (Figure 2DE); at a stringent enrichment p-value threshold of 10^-9^, lost enhancers are not enriched for any process while gained enhancers are enriched for 17 functions (Figure 2DE). As expected, conserved enhancers, which constitute the majority (72%) of all enhancers considered, are enriched for a large range of CNS developmental processes (Figure S5A and Table S3). Finally, we found that CNS related GWAS traits (Table S4-6) are enriched among *de novo* gained enhancers compared to conserved and lost enhancers (Figure 2F), suggesting an essential role of *de novo* gained enhancers in establishing cognitive traits.

**Figure 2.**
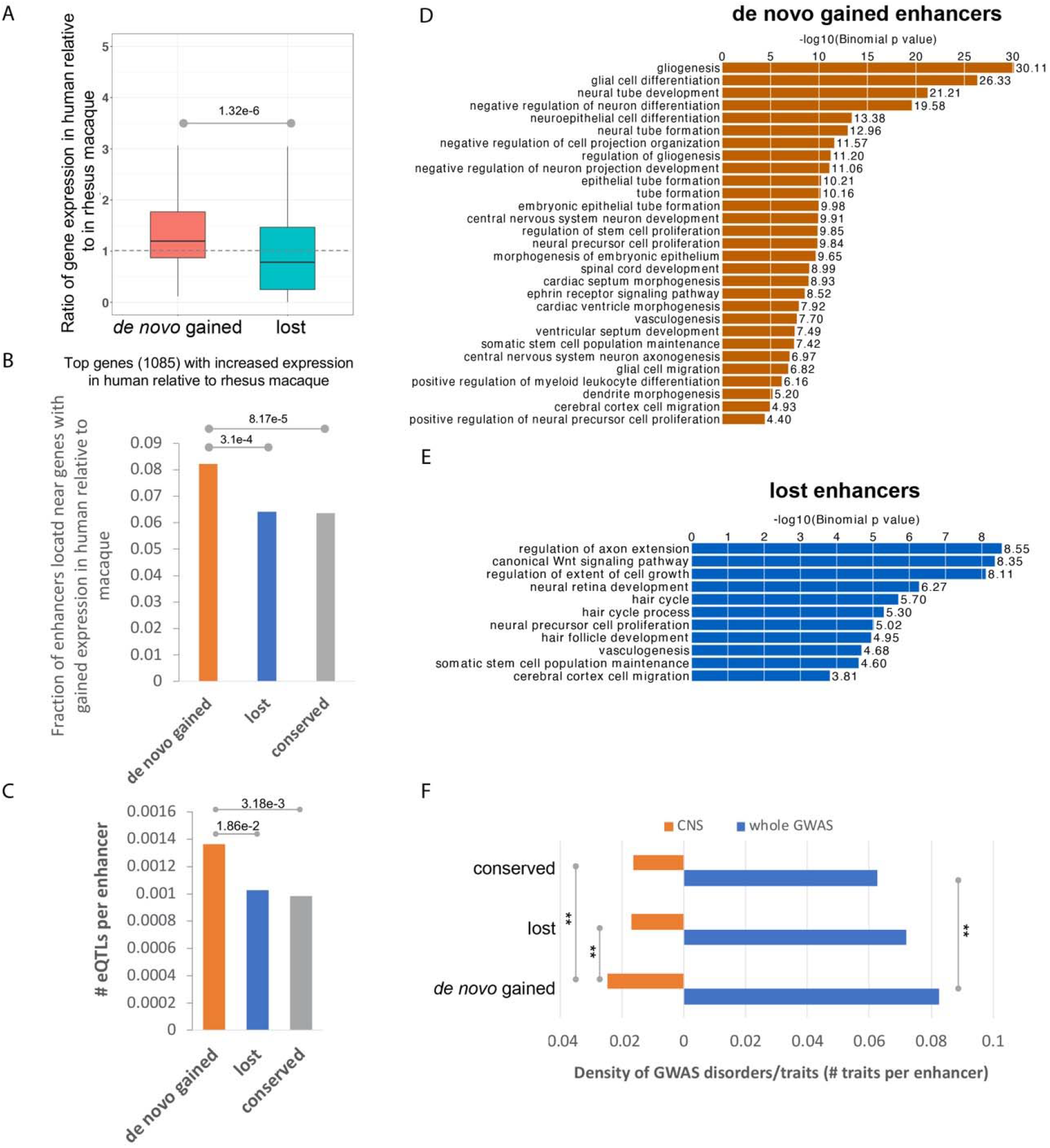
*De novo* gained enhancers are associated with essential biological pathways. (A) The expression level of genes near the *de novo* gained enhancers is increased. (B) Gained enhancers are enriched near the genes that are mostly highly expressed in humans as compared to rhesus macaque. (C) Average number of eQTLs per enhancer. (D) Biological processes that are associated with gained enhancers based on whole-genome region enrichment analysis performed using the GREAT tool (McLean et al. 2010). E) Biological processes that are associated with lost enhancers based on GREAT whole-genome region enrichment. F) The CNS related GWAS traits are enriched in the gained enhancers compared to both lost and conserved enhancers.

We further observed that, relative to conserved enhancers, *de novo* gained and lost enhancers are significantly enriched near genes that are specifically expressed in the embryonic neocortex (8 pcw), but not adult brain (Figure 3A, Methods), implicating them specifically in brain development. To fine map gained and lost enhancer activities to specific cell types of the developing human brain, we leveraged the single-cell transcriptomic data of developing human neocortex during mid-gestation (Polioudakis et al. 2019). Among the 16 transcriptionally distinct cell types/states (Figure 3B), *de novo* gained enhancers are primarily enriched near the genes specifically expressed in progenitor cells including radial glia (oRG, vRG), cycling progenitors in G2/M phase (PgG2M) and S phase (PgS), intermediate progenitors (IP), as well as interneurons (InCGE and InMGE), which connect different brain regions and are involved in cell/axon migration (Figure 3C and Figure S6). Although lost enhancers are enriched near genes specifically expressed in excitatory neurons (excitatory deep layers ExDp1 and ExDp2, maturing excitatory neurons ExM, ExM-u and migrating excitatory neurons ExN), *de novo* gained enhancers also exhibited a comparable level of enrichment in the same loci, thus arguing for compensatory impact on either the target gene expression or the phenotypic change to a large extent. Thus, the unique enrichment of *de novo* gained enhancers in the progenitor cells and interneurons might have contributed to the expansion of cortical surface and to an increased complexity of connections in the human cerebral neocortex, both of which together underpin the advanced cognition in humans. As such, in the following, we focus specifically on the *de novo* gained enhancers and investigate their emergence and functional consequences.

**Figure 3.**
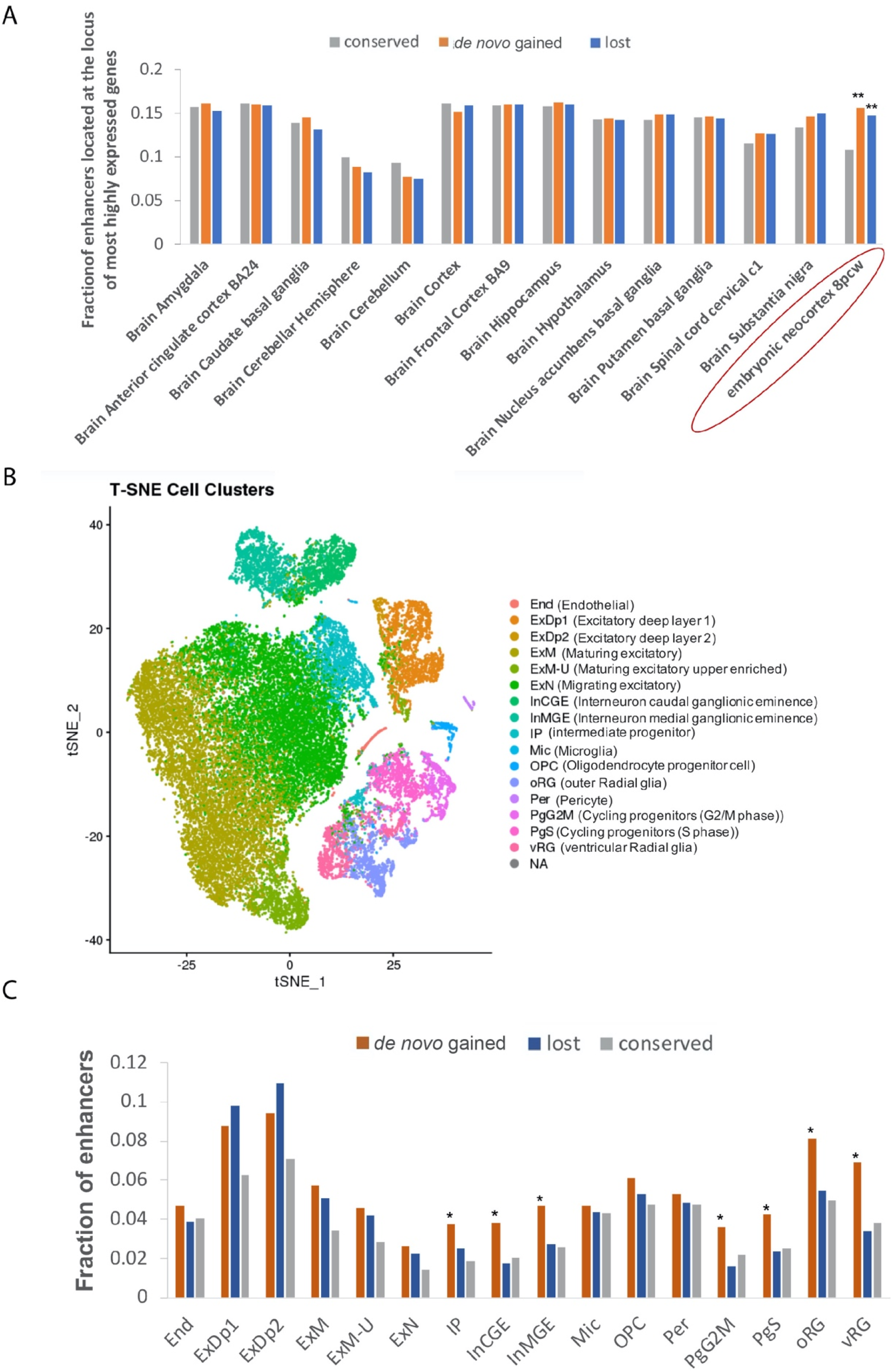
The *de novo* gained enhancers are enriched in the progenitor cells and interneurons. A) The *de novo* gained enhancers are significantly enriched in the most highly expressed genes of embryonic human neocortex but no other adult brain regions. ** indicates Fisher’s exact test P-value < 1e-3. B) Scatterplot visualization of cells after principal-component analysis and t-distributed stochastic neighbor embedding (tSNE), colored by Seurat clustering and annotated by major cell types. C) Fraction of enhancers near genes that are most highly expressed in all the cell clusters.

### A single essential mutation is often sufficient to create a human neocortical enhancer

To investigate the extent to which the enhancer gains could be explained by single-nucleotide mutations and to identify the minimal number of mutations needed to activate a neutral DNA sequence, we first compared the number of human-macaque mutations in *de novo* gained and conserved enhancers. The number of human-macaque mutations in *de novo* gained and conserved enhancers are comparable -- ~50 in a 1 kb enhancer (Figure S7). Recall that our DLM is trained to distinguish fetal brain enhancers from accessible non-fetal-brain-enhancer regions and not necessarily to assess the effect of single nucleotide changes. Therefore, we first performed a series of analyses to ensure that the DLM score (i) tracks enhancer activity and (ii) can accurately predict allele specific effects on H3K27ac signals (Supplementary Results 2). To identify critical mutations, we applied our DLM to prioritize human-macaque mutations in *de novo* gained enhancers based on the mutations’ impact on enhancer activity by iteratively introducing them into the potentially inactive macaque sequence orthologous to human CS23 enhancers. We were thus able to assess the minimal number of mutations capable of activating an enhancer (Methods). Even though only ~1.8% of all mutations in *de novo* gained enhancers are independently able to activate an enhancer (we call these essential mutations), ~40% of the *de novo* gained enhancers contain at least one essential mutation (Figure 4A). As expected, the smaller the minimal number of mutations needed to create an enhancer, the larger is their individual impact as per the DLM (Figure S8). To validate the impact of essential mutations on enhancer activity, we assessed their allelic imbalance of H3K27ac reads at the heterozygous sites. We hypothesized that the human reference allele at essential positions should exhibit larger H3K27ac read coverage than the macaque reference allele (Methods). Indeed, compared to three other groups of mutations/SNPs as controls, essential mutation positions are significantly associated with imbalance of H3K72ac reads coverage with the human reference allele (Figure 4B). This result strongly supports a causal link between the essential mutations and enhancer gain.

**Figure 4.**
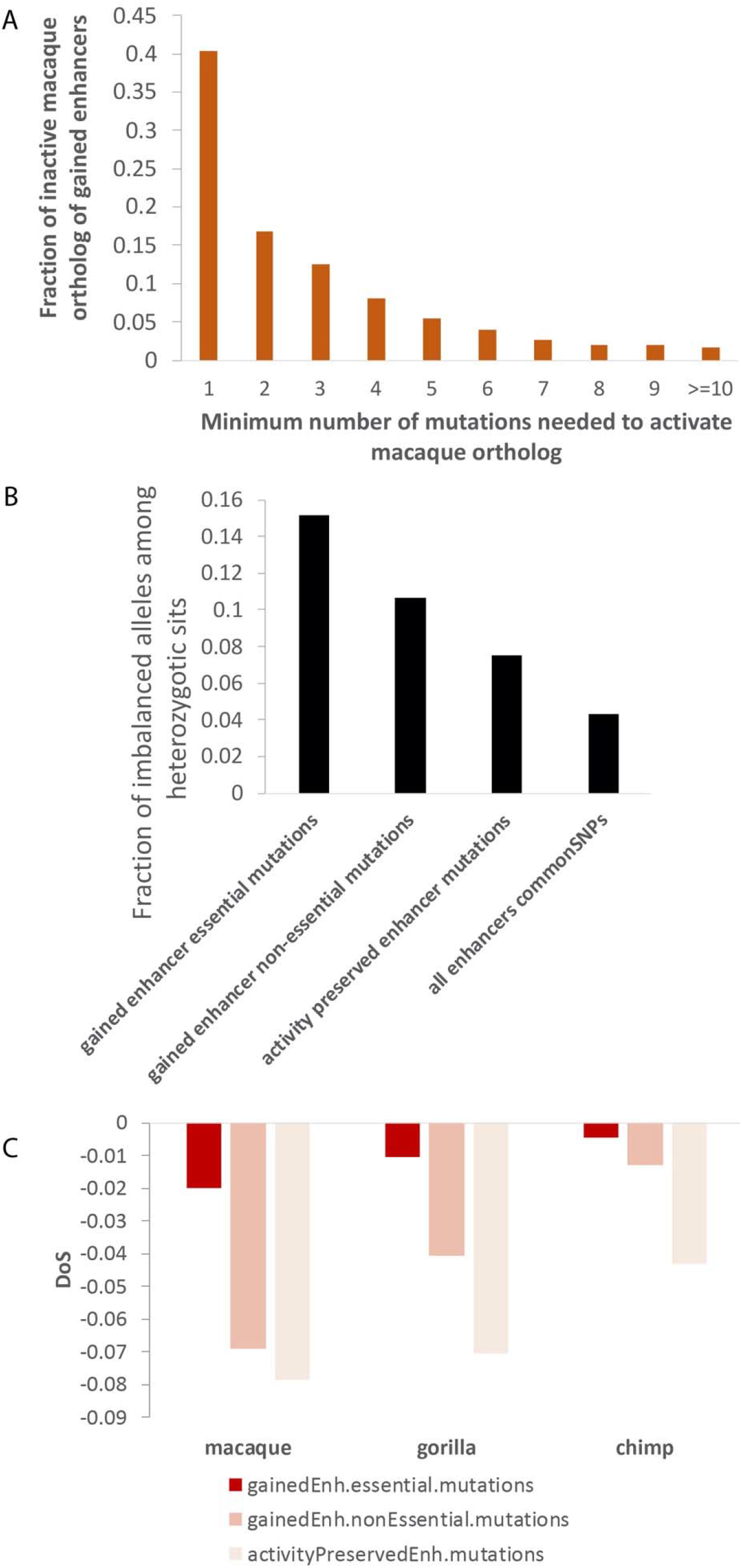
Essential mutations show larger impact on enhancer activity. A) Fraction of de novo gained enhancers that could be activated by specific number of mutations. B) Fraction of mutation/SNP sites that are in allelic imbalance. C) DoS score of the mutated sites, using macaque, gorilla, and chimp as comparison species.

We next examined the evolutionary constraints on essential mutations by applying the direction of selection (*DoS*) (Stoletzki and Eyre-Walker 2011) test, which is a refinement of McDonald-Kreitman (MK) test (Stoletzki and Eyre-Walker 2011), to measure the direction and degree of departure from neutral selection (Methods). *DoS* test is applied to a pair of species and a positive and negative *DoS* indicate positive and negative selection respectively. We estimated the *DoS* values for three sets of mutations -- essential mutations, non-essential mutations in *de novo* gained enhancers, and mutations within activity preserved enhancers (Methods) -- comparing human with macaque, gorilla, and chimp. As shown in Figure 4C, compared to other mutation classes, essential mutations have the highest *DoS* values, consistent with a relaxed negative selection, or potentially a subset of sites being under positive selection, both of which manifest as accelerated evolutionary rate (Cai and Petrov 2010; Hunt et al. 2011; Calderoni et al. 2016; Persi et al. 2016; Liu and Robinson-Rechavi 2018).

### Essential mutations are associated with cognition and neurodevelopmental disorders

Given our observation that the essential mutations are causally linked to enhancer activity in the embryonic neocortex, we assessed whether the essential mutations are preferentially associated with CNS-related GWAS traits (Methods). Indeed, we observed a ~2-fold enrichment of CNS related traits at the essential mutation positions as compared to non-essential mutation sites (Figure 5A, Table S7-8). Specifically, 7 out of 28 GWAS traits overlapping essential mutations are CNS related, and more importantly, 6 of those are associated with cognition (Table S7). We further investigated three such cases where the nearest genes are protein-coding genes with available expression data at approximate developmental stages (Zhu et al. 2018).

**Figure 5.**
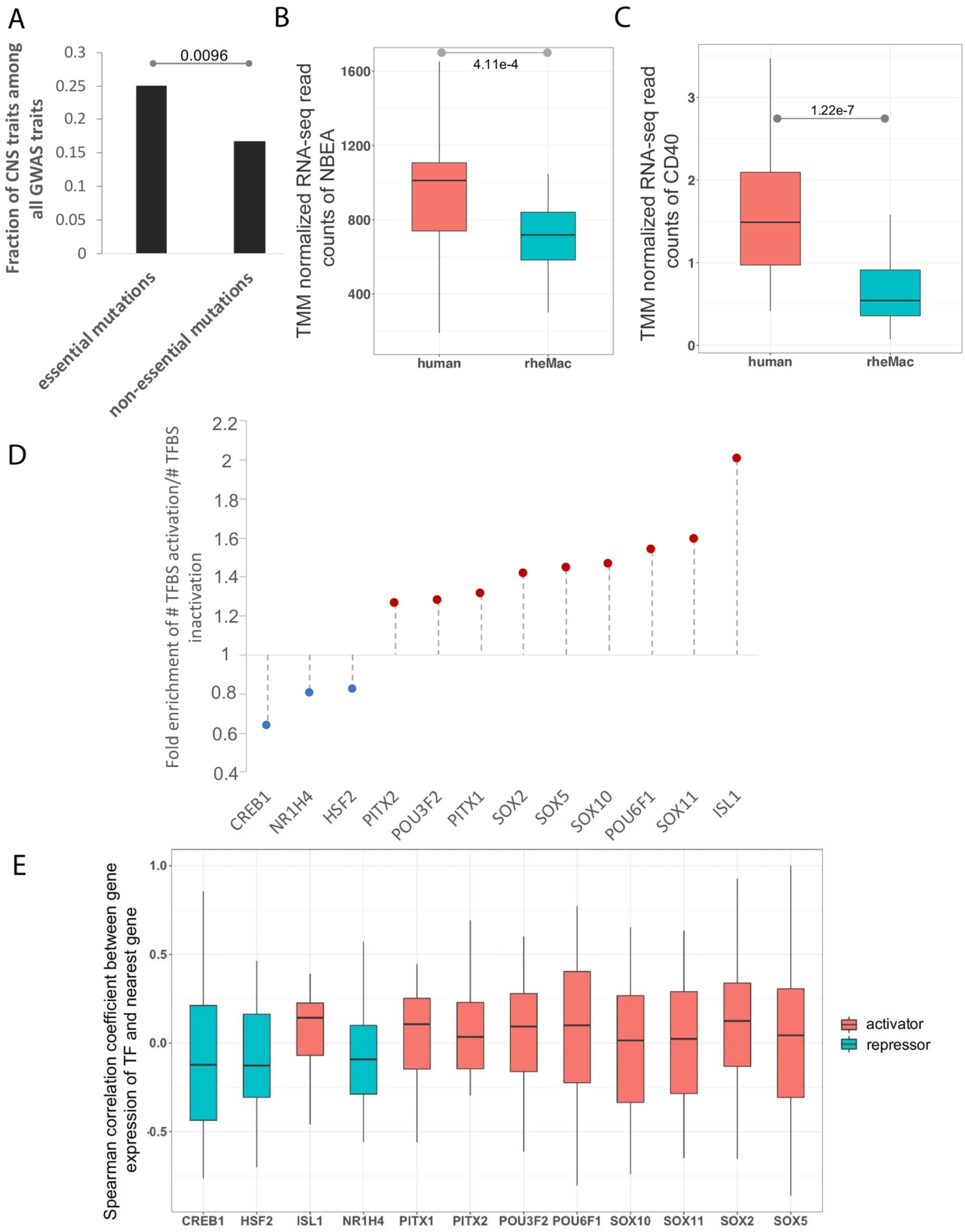
Essential mutations are associated with cognition related traits and tend to create binding sites of activators. A) Fraction of GWAS traits at the mutation sites which are CNS related. B) Comparison of TMM normalized expression of NBEA between embryonic human and rhesus macaque individuals. P-values are based on the Wilcoxon test. C) Comparison of TMM normalized expression of CD40 between embryonic human and rhesus macaque individuals. P-values are based on the Wilcoxon test. D) Enrichment of ratio of binding site gain to loss caused by essential mutations overlapping enriched TFBSs as compared to those caused by common SNPs. E) Spearman correlation coefficient of expression between the cognate TF of essential mutation and its nearest gene.

One essential mutation site coinciding with the common SNP rs9574096 is tightly linked to the tag SNP (rs9574095; correlation = 0.93) associated with the trait “Mathematical ability”. Both variants are located in the intronic region of the gene neurobeachin (NBEA), which is an autism-linked gene that fine-tunes signals at neuronal junctions (Nuytens et al. 2013). Mice missing one copy of NBEA show autism-like behavior (Nuytens et al. 2013). We found that NBEA exhibits a significantly higher embryonic neocortex expression in human compared to macaque at a similar early developmental stage (Zhu et al. 2018) (Figure 5B). Interestingly, the macaque allele A appears to be bound by another autism risk transcription factor, RFX3 (Harris et al. 2021), whereas the human allele T does not (Methods), suggesting a loss of RFX3 binding resulting in an increased enhancer activity and NBEA gene expression. Consistently, RFX3 expression is negatively correlated with that of NBEA in the embryonic neocortex across human and macaque individuals (Spearman rho = −0.26). In addition, NBEA is specifically expressed in sub-brain regions including excitatory neurons (ExDp1, ExDp2, ExM, ExM-U) and interneurons (InMGE) (Polioudakis et al. 2019), suggesting a link between these sub-brain regions and autism.

Other two essential mutation positions coincide with two common SNPs rs747759 and rs1535043, both of which are in perfect LD with each other. Notably, rs747759 is the tag SNP of the GWAS trait “Neuroticism”. The nearest gene of the two SNPs is CD40, which again displays a much higher expression in humans as compared to macaque (Figure 5C). CD40 is a major regulator of dendrite growth and elaboration in the developing brain (Carriba and Davies 2017) and contributes to synaptic degeneration in Alzheimer’s disease (AD) (Ye et al. 2019), which may have developmental origins (Arendt et al. 2017). The human allele T at the tag SNP rs747759 either causes a potential binding site gain of NFYA or a potential binding site loss of NHLH1 (Table S9). NFYA is an AD associated gene (Leslie et al. 2014; Nazarian et al. 2018; Nazarian et al. 2019). On the other hand, NHLH1 is known to play important roles in neuronal and glial differentiation and maturation (Dennis et al. 2019). However, the chance for NHLH1 to be a repressor of CD40 is dampened by their strong positive correlation of gene expression across human and macaque individuals (Spearman rho = 0.58). By contrast, NFYA expression is positively correlated with CD40 expression (Spearman rho = 0.29). At rs1535043, the human allele T is associated with the gain of an EHF binding site. However, its links with CNS traits are unclear.

Together, these results suggest a link between essential mutations in *de novo* gained enhancers and cognition-related traits as well as neurodevelopmental disorders in humans.

### Essential mutations tend to create binding sites of activating transcription factors

Next, we investigated the relative prevalence and importance of binding site gain versus loss in the *de novo* gained enhancers. Toward this, we focused on the TFs whose binding sites are enriched in the *de novo* gained enhancers compared to the conserved ones (using both human and macaque sequences to avoid allelic bias) (Table S10) and quantified the global tendency of essential mutations to lead to binding site gain versus loss (Methods). Overall, we observed that 9 TFs including POU3F2, PITX2, PITX1, SOX2, SOX5, SOX10, POU6F1, SOX11, and ISL1 tend to gain binding sites mediated by essential mutations in human (Figure 5D), suggesting an activator role of these TFs. Conversely, three TFs, CREB1, HSF2 and NR1H4, are more likely to lose their binding sites (Figure 5D), suggesting potentially repressive roles. Moreover, the overall positive or negative correlation of gene expression between these putative cognate TFs of the essential mutations and their nearest genes further validates their activator or repressor roles, respectively (Figure 5E). In short, the *de novo* gained enhancers are more likely to be activated by the creation of binding sites of activators due to the essential mutations.

### *De novo* gained enhancers induce a potential human-specific TF regulatory network

Transcriptional programs driving cell state are governed by a core set of TFs (also called master regulators), that auto- and cross-regulate each other to maintain a robust cell state. The ensemble of core TFs and their regulatory loops constitutes core transcriptional regulatory circuitry (Hnisz et al. 2013; Hnisz et al. 2015; Saint-André et al. 2016). Interestingly, the genes near *de novo* gained enhancers are enriched for transcriptional regulators (Figure S5B). We hypothesized that the TFs regulated by the *de novo* gained enhancers form a core regulatory network in the human embryonic neocortex. Toward this, first, we identified 24 TF genes (Table S11) near *de novo* gained enhancers and performed a motif scan for each of the 14 TFs having a known binding motif among all enhancers near the 24 TF genes (Methods). We found that the majority of the 14 TF motifs are enriched in the *de novo* gained enhancers near TF genes compared to the conserved enhancers in the same loci (Figure S9), suggesting a core regulatory network formed by these TFs. Next, we established a putative regulatory relationship for each TF pair based on the enrichment of the density of one TF’s motif in the *de novo* gained enhancer near another TF, including autoregulation, using conserved enhancers associated with the 24 TFs as the background (Figure 6AB). The inferred links are supported by our observation that linked TF pairs tend to have correlated expressions, as compared to those which are not (Figure 6CD). Based on the number of TFs each TF regulates, POU3F2 is likely to be the master regulator, with PITX2, TBX20, and PITX1 playing critical roles (Figure 6E). Moreover, we found the essential mutations that create a binding site for the TFs at higher hierarchical levels have a larger impact on the enhancer activity according to the DLM (Figure 6F). Interestingly, the *de novo* non-coding mutations in Autism patients (Zhou et al. 2019) are specifically enriched in the set of *de novo* gained enhancers associated with TF activity (Figure 6G). Remarkably, the *de novo* Autism mutations within this subset of *de novo* gained enhancers are more likely to be essential, which alone can deactivate an enhancer, as compared to those other *de novo* gained and conserved enhancers (Fig 6H). Together, these results suggest that essential mutations and the resulting enhancer gains may have helped create a core transcriptional regulatory network, with POU3F2 in a central position, to mediate a novel gene expression program in the developing human neocortex, associated with cognitive traits.

**Figure 6.**
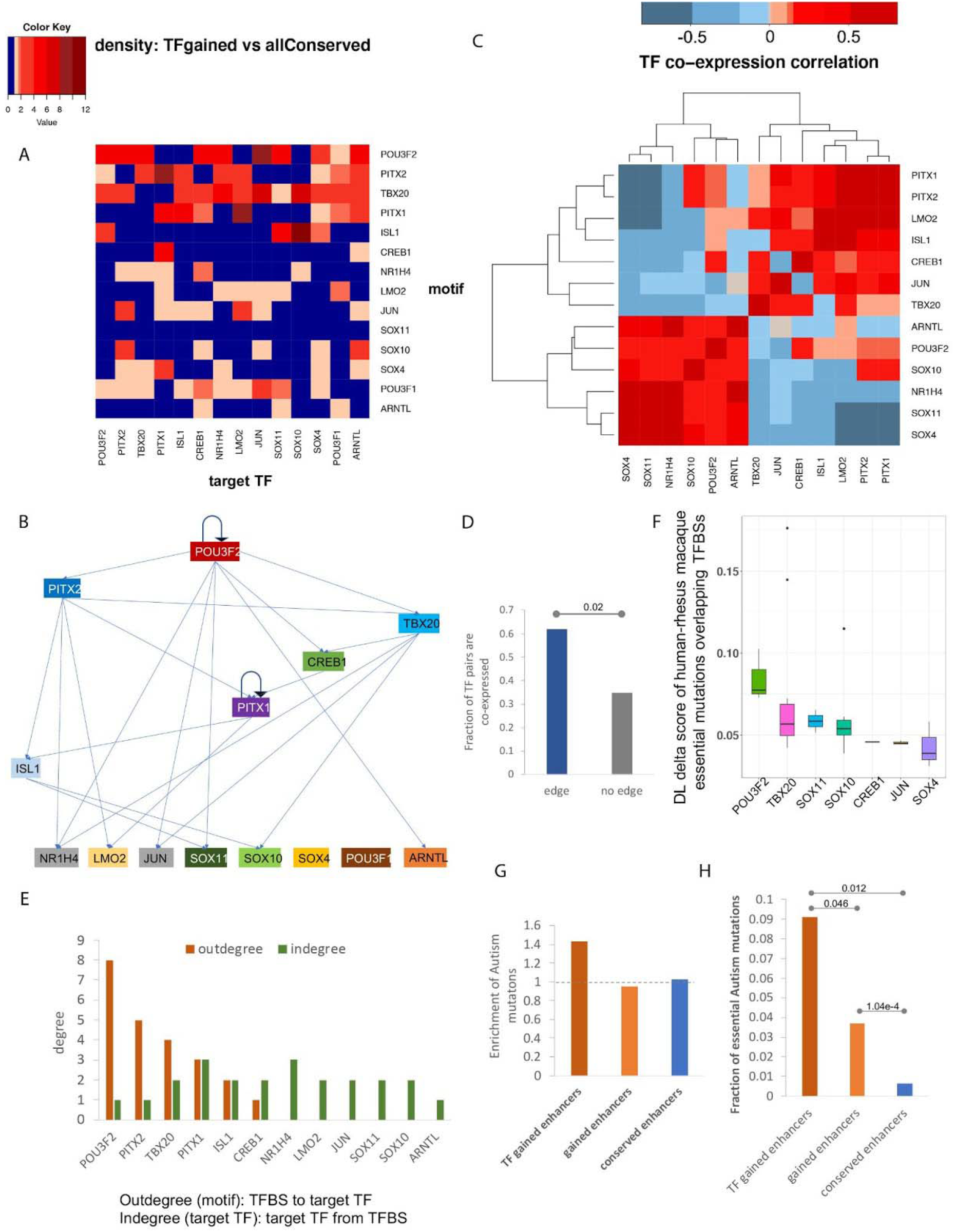
A hierarchical regulatory network of TFs induced by de *novo* gained enhancers. A) Density of TFBSs of the 14 TFs in the locus of the 14 TF genes. B) The inferred hierarchical structure of the 14 TFs. C) Spearman correlation coefficient of the 14 TF genes across the embryonic human and macaque individuals. D) Comparison of fraction of TF pairs that are co-expressed (Spearman correlation coefficient > 0.3) between the pairs with links and those without links. P-value is calculated using Fisher’s exact test. E) Out-degree and in-degree of each TFs. F) Distribution of DLM delta score caused by the essential mutations overlapping the 14 TFs. G) Fraction of Autism de novo mutations located within each set of enhancers normalized by the fraction of common SNPs falling into the same set of enhancers. H) Fraction of Autism *de novo* mutations within each set of enhancers, which are essential.

## Discussion

Higher cognition in humans is attributed to substantial expansion of the cortical surface and increased complexity of cortical connections during early development. Such phenotypic changes are likely to be mediated, in significant part, by changes in transcriptional regulation during brain development (Geschwind and Rakic 2013). Recent availability of genome sequencing and epigenomic data in the developing brain of humans and a close relative – rhesus macaque – has opened the possibility to probe key regulatory changes underlying the cognitive innovations in humans.

Here, we focused on one critical component of transcriptional regulation, namely, cis-regulatory enhancers. Our results suggest that single-nucleotide mutation in the human lineage, by creating binding sites for key TFs, may have induced novel enhancers which, mediated by a core regulatory network, involving POU3F2, PITX2, TBX20, and PITX1, underlie an increased expression in the developing neocortex of key genes involved in gliogenesis, neural tube development, and neuron differentiation. Further, analysis of scRNA-seq data from the developing human brain shows that the *de novo* gained enhancers are likely to be active specifically in the progenitor cells and interneurons, which notably, are thought to underlie the expansion of the cortical surface and connectivity in the human neocortex, respectively. Given that corticogenesis in human differ from other species mainly with respect to an increased duration of neurogenesis, increases in the number and diversity of progenitors, introduction of new connections among functional areas, and modification of neuronal migration (Schwartz et al. 1991; Rakic 2009), our results are highly suggestive of a mechanistic link between enhancer gains and higher cognition in humans. We also find that the *de novo* mutations in autistic individuals are especially enriched in the *de novo* gained enhancers associated with transcription activator activities, suggesting a shared basis between human cognition and autism.

Our *de novo* gained enhancers differ significantly from previously identified human gained enhancers (HGEs) (Reilly et al. 2015), both conceptually as well as in terms of various functional properties. Reilly et al. defined HGEs based on a comparative analysis of enhancer-associated epigenetic marks (H3K27ac and H3K4me2) in human with rhesus macaque and mouse. In other words, HGEs are enhancers with increased activity in human compared to both macaque and mouse. In sharp contrast, our “*de novo*” gained enhancers originate from presumably “neutral” non-coding sequence (i.e., without a detectable enhancer activity) in either macaque or the common ancestor of humans and macaques. In fact, HGEs are largely a subset of what we consider conserved enhancers in our study (85% CS23 HGEs overlap our conserved enhancers) and not *de novo* gained enhancers (only 11.9% overlap *de novo* gained enhancers). Notably, *de novo* gained enhancers exhibit weaker H3K27ac signals compared to the HGEs and conserved enhancers (Figure 7A), as they are largely activated by single-nucleotide mutations that potentially create binding sites of essential TFs in the developing brain (Figure 5DE). Therefore, it is not surprising that the *de novo* gained enhancers are more vulnerable to human substitutions that significantly alter enhancer activity (termed hSubs) according to Massively Parallel Reporter Assay (MPRA) targeted HGEs (Figure 7B, Methods) (Uebbing et al. 2021). Importantly, the *de novo* gained enhancers are more likely to turn on the expression of a gene in human compared to the HGEs (Figure 7C and Figure S10A), as the macaque counterpart of the human *de novo* gained enhancers are inactive in embryonic neocortex, whereas the macaque counterpart of HGEs is also an active enhancer, albeit relatively weaker. Therefore, all else being equal, the macaque orthologs of human genes associated with *de novo* gained enhancers are more likely to be silent.

**Figure 7.**
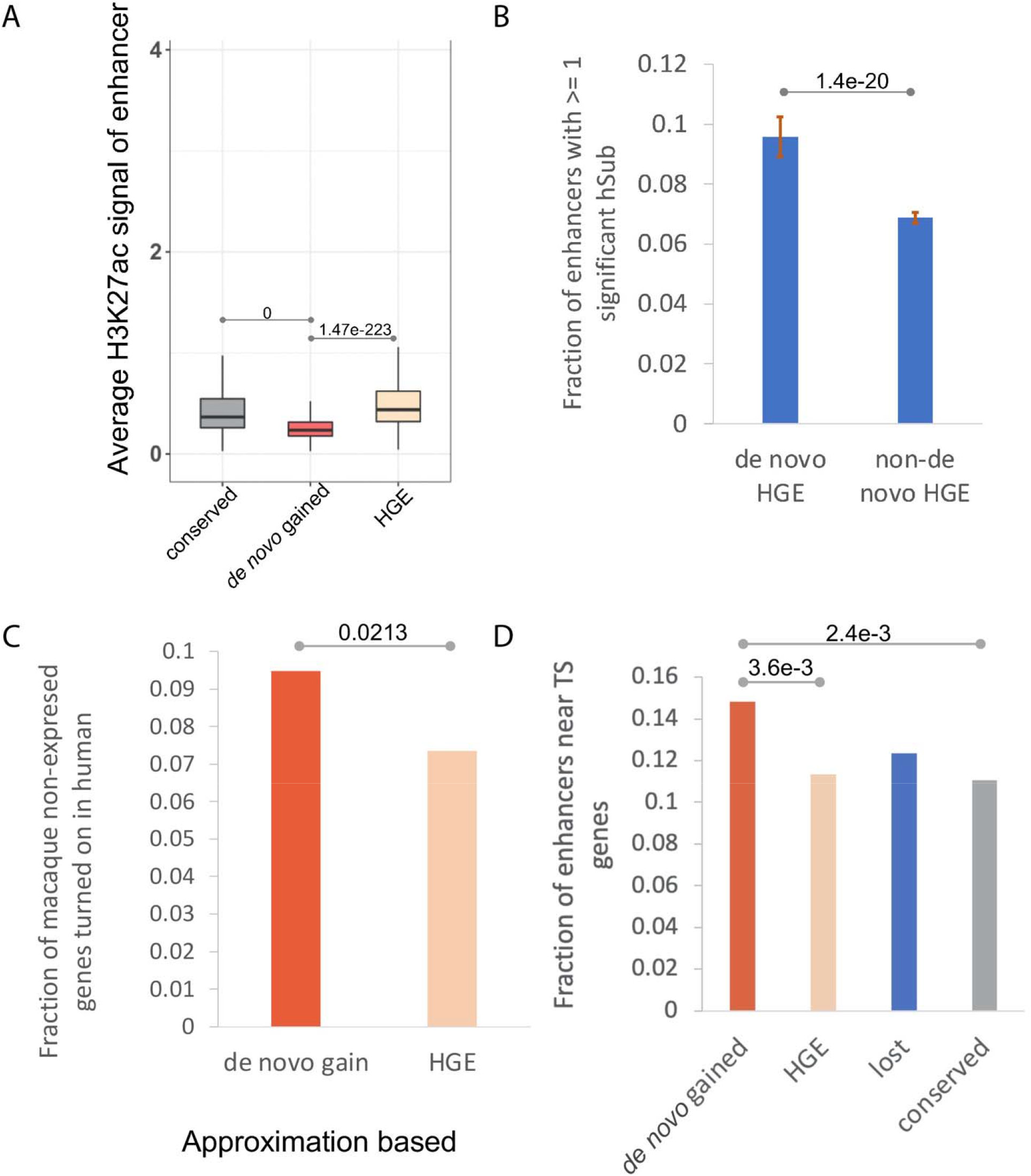
*de novo* gained enhancers versus HGEs. (A) *de novo* gained enhancers exhibit weaker enhander signal. (B) Fraction of enhancers with >= 1 significant hSubs. Bar plot shows the median and standard deviation of fraction of enhancers with at least one significant hSub by 90% Bootstrapping for 50 times. P-value is based on t-test. *De novo* HGEs refers to HGEs that overlap *de novo* gained enhancers, and non-*de novo* HGEs refers to CS23 HGEs that do not overlap *de novo* gained enhancers. (C) Fraction of enhancers located near genes whose RPKM < 1 in macaque and > 1 in human. (D) *de novo* gained enhancers are more likely to locate near tissue-specific (TS) genes.

In addition, the brain morphology related functions were reported to be associated with HGEs by the earlier study (specific functions in neuronal proliferation, migration, and cortical-map organization) (Reilly et al. 2015), which differs notably from our findings, which implicate human *de novo* gained enhancers specifically in human neocortex development. Furthermore, GO enrichment analysis based on either nearby genes (Figure S11) or genes linked via Hi-C contacts (Table S1 and Table S12) consistently shows that *de novo* gained enhancers are more likely to be associated with more tissue-specific functions of the developing human brain compared to HGEs (Supplementary Results 3). Indeed, *de novo* gained enhancers are more likely to reside near (Figure 7D) or at 3D contact positions (Figure S10B) with the most tissue-specific genes in embryonic neocortex (Table S13). As mentioned above, *de novo* gained enhancers exhibit weaker H3K27ac signals compared to the HGEs. Previous studies have implicated weaker enhancers to be specifically critical during development (Farley et al. 2015), further suggesting a link between *de novo* gained enhancers and brain development.

## Methods

### Data Availability

We downloaded the gene expression data in the prenatal neocortex of human and macaque (Zhu et al. 2018). For human, we chose the time-points at 8 p.c.w and 12 p.c.w; for macaque, we selected the approximately matching time-points at E60 and E82 (Table S14). The data is shared by the authors at http://evolution.psychencode.org/#. The single-cell transcriptomic data of developing human neocortex during mid-gestation (Polioudakis et al. 2019) is shared by the authors at http://solo.bmap.ucla.edu/shiny/webapp/. In addition to assigning enhancers to their nearby genes, we also used the Hi-C loops in the developing brain of human (Won et al. 2016) and macaque (Luo et al. 2021) to link enhancer to their gene targets. The CS23 HGEs were obtained from the study (Reilly et al. 2015). All the potential fetal brain enhancers (the merged ATAC-seq peaks from the germinal zone and cortical plate of the human developmental brain) were obtained from the study (de la Torre-Ubieta et al. 2018). The fetal brain eQTLs were obtained from the study (O’Brien et al. 2018).

### Embryonic neocortex enhancers in human and rhesus macaque

The H3K27ac peaks of both species were obtained from the previous study (Reilly et al. 2015). The enhancers were defined as H3K27ac peaks extended to 1 kb from its original center. Integrating wider sequence context is critical because sequence surrounding the variant position determines the regulatory properties of the variant, as in vivo TF binding depends upon sequence beyond traditionally defined motifs (Deplancke et al. 2016; Inukai et al. 2017). Enhancers overlapping promoters (including all alternative promoters) and promoters (intervals [-1000 bp, 1000 bp] surrounding the transcription start site) were removed from the enhancer set. Overall, we identified 32,201 human enhancers, and 43,997 macaque enhancers.

### A deep convolutional neural network model for enhancer prediction

We built a deep convolutional neural network to predict tissue-specific enhancer activity directly from the enhancer DNA sequence. The DLM comprises 5 convolution layers with 320, 320, 240, 240, and 480 kernels, respectively (Table S15). Higher-level convolution layers receive input from larger genomic ranges and are able to represent more complex patterns than the lower layers. The convolutional layers are followed by a fully connected layer with 180 neurons, integrating the information from the full length of 1,000 bp sequence. In total, the DLM has 3,631,401 trainable parameters. We used the Python library Keras version 2.4.0 (https://github.com/keras-team/keras) to implement our model.

The model was trained for each of the four temporal-spatial groups of enhancers (CS16, CS23, F2F, and F2O). The positive sets contain the human embryonic enhancers of each group. The Dnase I-hypersensitive sites (DHSs) profiles of non-CNS-related and non-embryonic tissues from Roadmap Epigenomics projects (Kundaje et al. 2015), which do not overlap the positive sets, were collected as the negative training set of the DL model. The reason we used DHS sites not overlapping embryonic neocortex H3K27ac peaks as negative control regions is that we aim to identify tissue-specific enhancers of embryonic neocortex, and DHS is a good representation of active chromatin. The fact that DHS in general overlaps H3K27ac makes it a stringent control, and in fact, our choice of DHS as the control is analogous to DeepSEA, which utilizes the genomic regions not overlapping the positive set and with at least one TF binding as the negative set, which broadly overlap with DHS regions.

Training and testing sets were split by chromosomes. Chromosome 8 and 9 were excluded from training to test prediction performances. Chromosome 6 was used as the validation set, and the rest of the autosomes were used for training. Each training sample consists of a 1,000-bp sequence (and their reverse complement) from the human GRCh37 (hg19) reference genome. Larger DL score of the genomic sequence corresponds to a higher propensity to be an active enhancer. The genomic sequence with DLM score >= 0.197 (FPR <= 0.1) are predicted to be active enhancers. We used the difference of the DLM score induced by a human-macaque single-nucleotide mutation to estimate its impact on enhancer activity.

Given a human (hg19) or macaque (rheMac2) enhancer, we used liftOver (Hinrichs et al. 2006) to identify their orthologs. Only the reciprocal counterparts with their lengths difference no more than 50 bp were considered to be ortholog pairs. For a human sequence with *n* mutations relative to its macaque ortholog, to score the impact of combinations of *m* (*m* < *n*) mutations on enhancer activity, all possible combinations of *m* (*n* choose *m*) human alleles at the human-macaque mutation sites were introduced to the macaque orthologs if the total number of combinations (*n* choose *m*) is no more than 10,000, otherwise, we randomly sample 10,000 combinations of m human alleles from the human-macaque mutation sites and introduce them to the macaque ortholog. The change of DL score caused by the set of introduced human mutations were used to estimate their impact on enhancer activity.

### Gain and loss of enhancers

Briefly, if a human enhancer having a high DLM score scored low both in macaque and in the common ancestor, and was not detected by H3K27ac in macaque, it was considered to be a *de novo* gain in humans (Figure 1A). Likewise, if a macaque enhancer having high DL score scored high in common ancestor, scored low in human and was undetectable by H3K27ac in human, it was considered a loss in human (Figure 1A). The enhancers that are detected by H3K27ac in both human and macaque, and scored highly in all three genomes were called conserved enhancers (Figure 1A).

### Normalization of gene expression data

We applied ‘tmm’ built-in normalization method of edgeR to normalize human and macaque embryonic neocortex gene expression and to remove differences across species and batch effects. To identify the most tissue-specific genes of human embryonic neocortex, the expression data of human individuals were averaged and quantile normalized together with the gene expression profile downloaded from GTEx. The top 2000 genes with the highest ratios of the human embryonic expression to the mean of the GTEx expression were identified as the most specifically highly expressed genes in human embryonic neocortex (Table S13).

### *De novo* single-nucleotide substitutions in autism spectrum disorder (ASD)

We obtained 127,141 *de novo* single-nucleotide mutations in ASD from a previous study (Zhou et al. 2019), which were identified from Simons Simplex Collection of whole-genome sequencing data for 1790 families that were available via the Simons Foundation Autism Research Initiative (SFARI).

### Functional enrichment analysis using GREAT and DAVID tools

To probe the potential functional roles of gained and lost enhancers we first tested for functional enrichment among genes near the enhancer loci using the online Genomic Regions Enrichment of Annotations Tool (GREAT) version 3.0.0 (McLean et al. 2010) using single nearest gene association rule with more strict settings than default. Specifically, the GO terms will be considered as enriched if it has at least 10 gene hits with FDR threshold set as 0.01. Two background options were used when using GREAT. Figure 2DE, Figure S5 and Figure S11 are based on enrichment against whole genome region. Next, we performed GO enrichment analysis using all potential fetal brain enhancers (the merged ATAC-seq peaks from the germinal zone and cortical plate of the human developmental brain) (de la Torre-Ubieta et al. 2018) as the background and obtained consistent observations (Figure S4AB and Figure S12AB). The exception is the conserved enhancers, which are not enriched for CNS related biological processes (Figure S12A). The tissue-specific signal of conserved enhancers is dampened, as expected, by using the fetal brain enhancers as the background, as the conserved enhancers constitute the majority of the fetal brain enhancers.

We also applied DAVID (Huang da et al. 2009b; Huang da et al. 2009a) to do functional enrichment of the genes with Hi-C loops to different sets of enhancers.

### Enrichment analysis of GWAS traits and eQTLs

The NHGRI-EBI GWAS Catalog (Buniello et al. 2019) was downloaded. To study the enrichment of a set of SNPs coinciding with CNS related GWAS traits, the tag SNPs were first expanded by linkage disequilibrium (LD) (r2 >⍰0.8, maximum distance of 500⍰kb) using Plink (http://pngu.mgh.harvard.edu/purcell_urcell/plink/; (Purcell et al. 2007)) with the following parameters:

‘--r2 --ld-window-kb 500 --ld-window-r2 0.8’

We overlapped the LD-expanded GWAS traits with the human-macaque mutation sites of the gained enhancers where the human alternative alleles are the same as the macaque reference alleles. The CNS-related GWAS traits are listed in Table S4-6. We then use the fraction of CNS-related traits among the total GWAS traits overlapping the essential mutations, as compared to that of the non-essential mutations to estimate the enrichment of CNS-related traits in the essential mutation positions (Figure 5A).

As for the overall enrichment of the CNS-related GWAS traits in the three sets of enhancers (Figure 2F), we used the density (average number of LD-expanded GWAS traits per enhancer) to estimate the enrichment. As the density of common SNPs in the three sets of enhancers (average number of SNPs per enhancer) is comparable (gained: 4.1, lost: 4.05, conserved: 4.6) and would not change the trend of the enrichment upon normalization, we did not normalize the GWAS density by SNP density.

As for the enrichment of the fetal brain eQTLs (O’Brien et al. 2018) in the three sets of enhancers, we first compared the density of eQTLs (average number of eQTLs per enhancer) in the three sets of enhancers (Figure 2C). Next, we normalized the fraction of eQTLs fallen within a set of enhancers by the fraction of common SNPs fallen within that set of enhancers (Figure S3).

### Identification of potential TFBSs in the *de novo* gained enhancers

To identify potential binding sites, we used FIMO (Bailey et al. 2009) to scan the profiles of binding sites for vertebrate TF motifs in Jaspar (Mathelier et al. 2014), CIS-BP (Weirauch et al. 2014), SwissRegulon (Pachkov et al. 2007), HOCOMOCO (Kulakovskiy et al. 2016), and UniPROBE (Hume et al. 2015) databases, along the enhancer sequences. We identified motif-specific thresholds to limit the false discovery rate to no more than five false positives in 10⍰kb of sequence, by scanning each motif on random genomic sequences using FIMO (Bailey et al. 2009). Enrichment of a motif in *de novo* gained (foreground) relative to conserved (background) enhancers were ascertained using Fisher’s exact test. The occurrence of a particular TFBS in the set of *de novo*-gained/conserved sequences was normalized by the total number of *de novo*-gained/conserved regions.

However, when identifying TFs whose motifs are enriched in *de novo* gained enhancers relative to conserved enhancers, we included both the human and the macaque ortholog sequences, to avoid allelic bias in our following analysis of activation/repression of enhancers by single-nucleotide mutations. Next, we assessed whether a mutation (in a *de novo* gained enhancer) creates a binding site of a potential activator or disrupts binding of a potential repressor, we estimated, for each enriched TF, the ratio of binding site gain to loss caused by essential mutations within *de novo* gained enhancers relative to the same ratio caused by common SNPs. If the gain/loss (loss/gain, respectively) ratio caused by essential mutations was greater than 1.2-fold that for common SNPs, the TF was considered activator (repressor, respectively).

### Identification of allelic imbalance in H3K27ac data

We used BWA (Li and Durbin 2010) to map two replicates of CS23 H3K27ac data (Reilly et al. 2015) to hg19 human reference sequence. At the mutation/SNP sites, the H3K27ac reads were extracted using BaalChIP (de Santiago et al. 2017). Allelic counts over heterozygous sites of the two replicates were merged, and variants that had at least 6 reads were further processed for allele specific enhancer activity analysis with Binomial test. We use the heterozygous sites within the activity preserved enhancers (the ratio between human and macaque H3K27ac signal is no more than 1.2) as the background. For a heterozygous site, if the ratio of reads number of the human allele to that of the macaque allele is over 1.3 and the Binomial p-value <= 1e-3, the position is considered to have allelic imbalance.

### Single-cell clustering and visualization

Clustering was performed using Seurat (v2.3.4) (Stuart et al. 2019). Read depth normalized expression values were mean centered and variance scaled for each gene, and the effects of number of UMI (sequencing depth), donor, and library preparation batch were removed using a linear model with Seurat (‘ScaleData ‘function). Highly variable genes were then identified and used for the subsequent analysis (Seurat ‘MeanVarPlot ‘function). Briefly, average expression and dispersion are calculated for each gene, genes are placed into bins, and then a z-score for dispersion within each bin is determined. Principal component analysis (PCA) was then used to reduce dimensionality of the dataset to the top 13 PCs (Seurat ‘RunPCA ‘function). Clustering was then performed using graph-based clustering implemented by Seurat (‘FindClusters ‘ function). Cell clusters with fewer than 30 cells were omitted from further analysis. Clusters were annotated using the Seurat function ‘group.by’.

For visualization, t-distributed stochastic neighbor embedding (tSNE) coordinates were calculated in PCA space, independent of the clustering, using Seurat (‘RunTSNE ‘function). tSNE plots were then colored by the cluster assignments derived above, gene expression values, or other features of interest. Gene expression values are mean centered and variance scaled unless otherwise noted.

### Direction of selection test

The *DoS* test was designed to measure the direction and extent of departure from neutral selection based on the difference between the proportion of substitution and polymorphism in the selective sites. *DoS* is positive when there is evidence of adaptive evolution, is zero if there is only neutral evolution, and is negative when there are slightly deleterious mutations segregating (Stoletzki and Eyre-Walker 2011). Here, we used the mutated four-fold degenerate sites as the background to measure the selection on the mutations within *de novo* gained enhancers (formula 1). Note that all sites in our three mutational site classes are, by design, mutated in human relative to macaque. Therefore, to avoid ascertainment bias, we uniformly applied the same criteria of human-macaque mutation to select a subset of all fourfold degenerate sites.

Let, *n* represent the ‘non-synoymous ‘sites, i.e. the essential or non-essential mutations within the *de novo* gained enhancers. *S* represents the ‘synonymous ‘sites, i.e. the mutated four-fold degenerate sites. *D* means ‘diverged ‘sites, i.e. mutations (or substitutions) that are fixed in the human populations, and *P* means ‘polymorphic ‘sites, i.e. both the ancestor allele and the mutations are preserved in the human populations (Table 1).

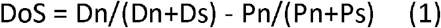

**Table 1.**
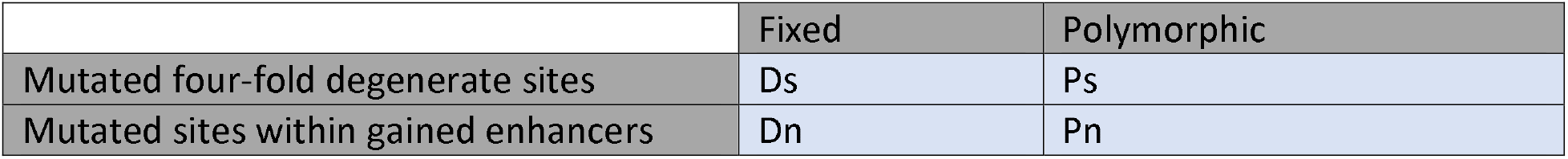
Contingency table of number of fixed mutations and polymorphic mutations at the foreground and background sites.

### Comparing *de novo* gained enhancers and HGEs using MPRA data

Overlapping the significant human substitutions (relative to chimp, termed hSubs) from a MPRA targeted HGEs (Uebbing et al. 2021) with the *de novo* gained enhancers, we found that 141 *de novo* gained enhancers overlapping HGEs (dubbed *de novo* HGEs) were tested by this assay. In total, 14 of the 141 (10%) *de novo* HGEs harbor at least one hSub. For the 1,019 CS23 HGEs that do not overlap *de novo* gained enhancers (dubbed non-*de novo* HGEs), 74 (7%) HGEs have at least one hSubs (Figure 7B). We applied 90% bootstrapping 50 times to estimate the statistical significance of the difference between the two fractions (Figure 7B).

## Acknowledgement

This work utilized the computational resources of the NIH HPC Biowulf cluster and was supported by the Intramural Research Program of the National Cancer Institute, Center for Cancer Research, and National Library of Medicine, NIH. We would like to thank Di Huang, Vishaka Gopalan, and Arashdeep Singh for their feedback. We would also to thank James Noonan and Jian Zhou for nicely providing supplementary materials of their works.

## Supplementary Materials

### Supplementary Results

#### 1. Performance and further validation of DLM of embryonic neocortex enhancers

Here we provide benchmarking and comparison of our enhancer model with DeepSEA (Zhou and Troyanskaya 2015).

To directly compare our model performance with DeepSEA, we applied our model to the training and testing H3K27ac data sets used by DeepSEA. Our model achieved a very similar (although slightly higher) accuracy (both auROC and auPRC) compared to DeepSEA across multiple datasets (Figure S13BC).

We have shown that the human embryonic neocortex DLM can accurately estimate the enhancer activity (independently) in macaque from its genomic sequence (Fig 2A). To further validate our model, we applied the model trained on the human embryonic neocortex (CS23) enhancers (H3K27ac peaks) and tested it on the mouse embryonic neocortex enhancers (H3K27ac peaks) (Reilly *et al* 2015, PMID:25745175), using random genomic regions (due to a lack of available multi-tissue DHS profile) that do not overlap H3K27ac peaks as the negative testing set. Even for this more distant species, the model achieves an auROC of 0.9 at e11 (Figure S13D).

#### 2. DLM can accurately predict allele specific effects on histone marks H3K27ac

Our DLM is trained to distinguish enhancer region from non-enhancer regions in a specific context. However, its application to identify *de novo* enhancer gains driven by single nucleotide mutations requires the DLM score to be sensitive to single nucleotide changes. We performed additional analyses to ensure that DLM score indeed (i) represent the enhancer activity level, and (ii) is sensitive to single nucleotide changes.

First, we computed the direct correlation between the predicted DLM score (DL score) of the enhancers and the log of their average H3K27ac signal intensity. We observed a significant positive correlation between the two (correlation = 0.4, empirical p-value = 3.18e-6).

Next, DeepSEA was shown to work well in identifying variants at loci that affect histone signals (hQTLs of H3K27ac or H3K4me3) (Zhou and Troyanskaya 2015). As our approach is very similar to DeepSEA (just a different neural net architecture), and we aim to identify variants that create enhancers, we trained our model on H3K27ac peaks in a lymphoblastoid cell line, GM12878, and applied it to predict the same set of hQTLs of H3K27ac in lymphoblastoid cell lines (McVicker, G. et al. Science 342, 747-749 (2013)) as did DeepSEA. Our model shows similar accuracy as DeepSEA (Figure S14).

To further show the ability of our DLM to accurately predict chromatin features from sequence with single-nucleotide sensitivity, we applied our CS23 model to evaluate the 2,578 allelically imbalanced SNPs within the CS23 H3K27ac peaks, which were identified using the R-package BaalChIP (de Santiago et al. 2017). Our model makes similarly accurate predictions on this set of SNPs as well (Figure S15).

#### 3. Using Hi-C loops to link enhancers to their potential target genes

In the main result sections, we opted to use proximity as the criterion to identify the enhancer-associated gene for several reasons. First, the available human Hi-C contacts (Won et al. 2016) are very sparse: only 23% of human embryonic neocortex enhancers are covered. The 3D contacts in macaque (Luo et al. 2021) are even sparser, where 8,399 and 15,048 loops were identified in the germinal zone and cortical plate, covering only 2.68% of total macaque enhancers. Second, in the study of ‘Activity-by-Contact model’ (Fulco et al. 2019), based on a small number of experiments, the authors concluded that it is rare for an enhancer to skip the nearest gene (Fulco et al. 2019). Finally, for the enhancers included in Hi-C loops, around 60% of *de novo* gained enhancers contact their nearest genes, and more than 50% of both lost and conserved enhancers are in contact with their nearest genes (Figure S16), suggesting that our findings based on the nearest genes are robust.

Nevertheless, we examined the results when the enhancers were mapped to their putative targets based on Hi-C loops. The findings based on the Hi-C loops are consistent with the ones based on the proximity rule. For example, the *de novo* gained enhancers tend to associate with an increase in the expression of their target gene, whereas the lost enhancers show the reverse trend (Figure 2A and Figure S2). Enhancers are more likely to regulate the tissue-specific genes of embryonic neocortex either based on proximation rule (Figure 7B) or Hi-C contacts (Figure S10B). In addition, using either gene proximation rule (Figure 7C) or Hi-C contact (Figure S10A), we observed that *de novo* gained enhancers are more likely to turn on gene expression compared to HGEs.

## Supplementary Figures

**Figure S1.**
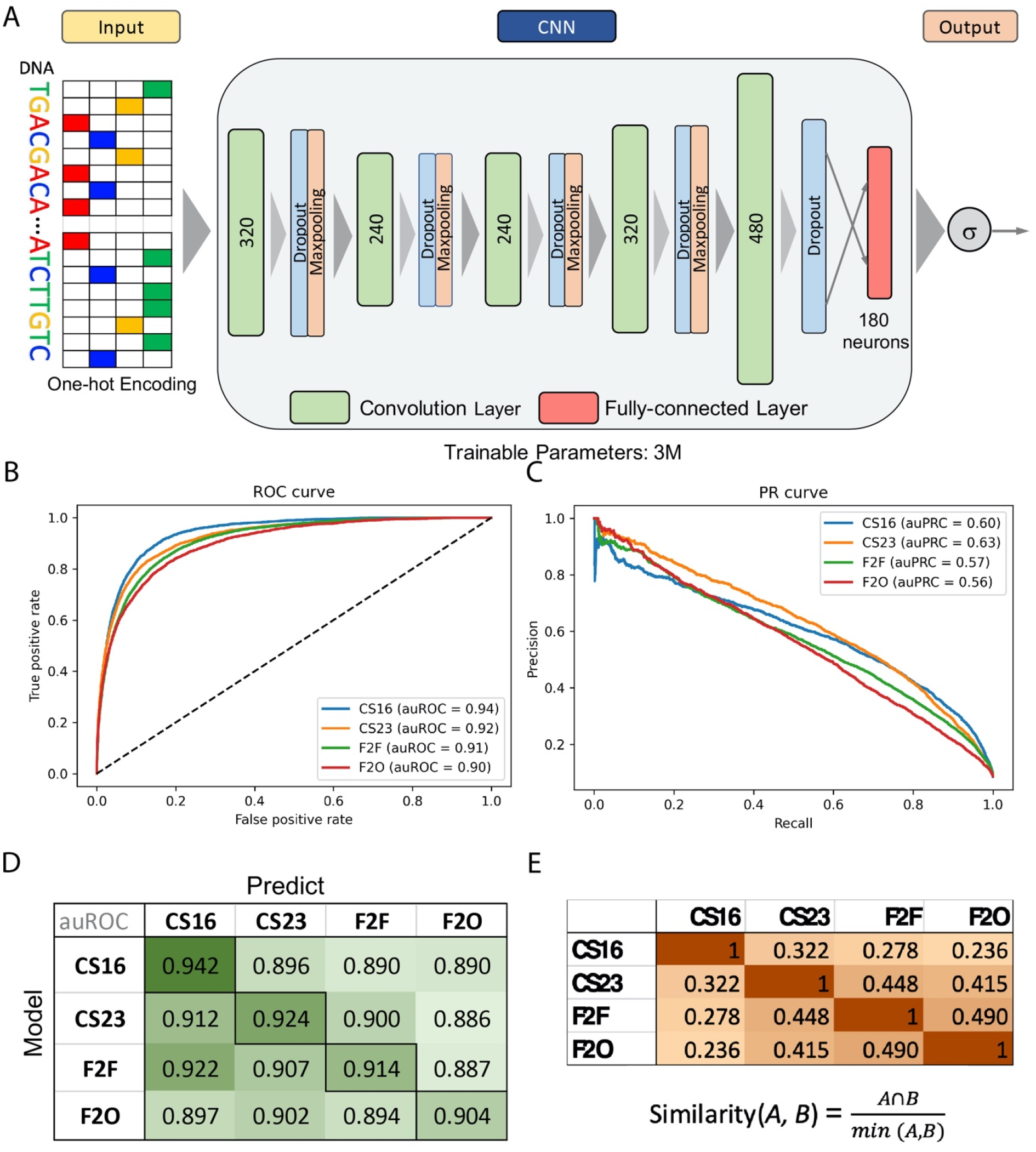
Deep learning model of human embryonic neocortex enhancers used to score enhancer activity. A) Structure of the deep convolutional model. The number within each convolutional layer indicates the number of kernels. B) ROC curve of the model. C) PR curve of the model. D) Model performance across four stages. E) Similarity between enhancer sets across stages.

**Figure S2.**
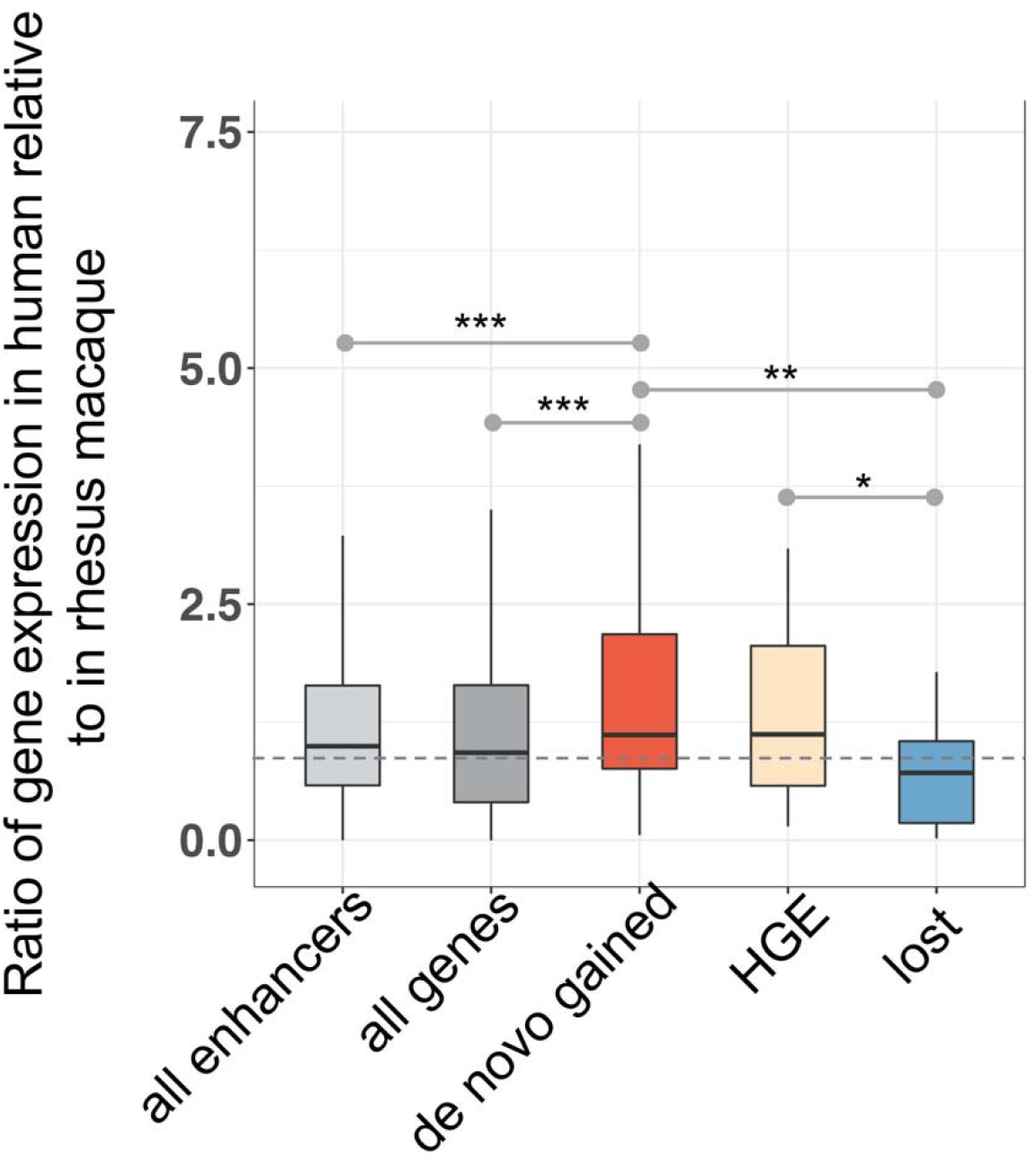
The expression level of genes with Hi-C loops to the *de novo* gained enhancers is increased, and so is the previously published enhancers that increase activity in human (HGEs, PMID: 25745175). By contrast, the genes in contact with the lost enhancers show the reverse trend, “all enhancers” refer to the genes link to all enhancers. *Wilcoxon p-value <= 0.01. ** Wilcoxon p-value <= 1e-3. *** Wilcoxon p-value <= 1e-5.

**Figure S3.**
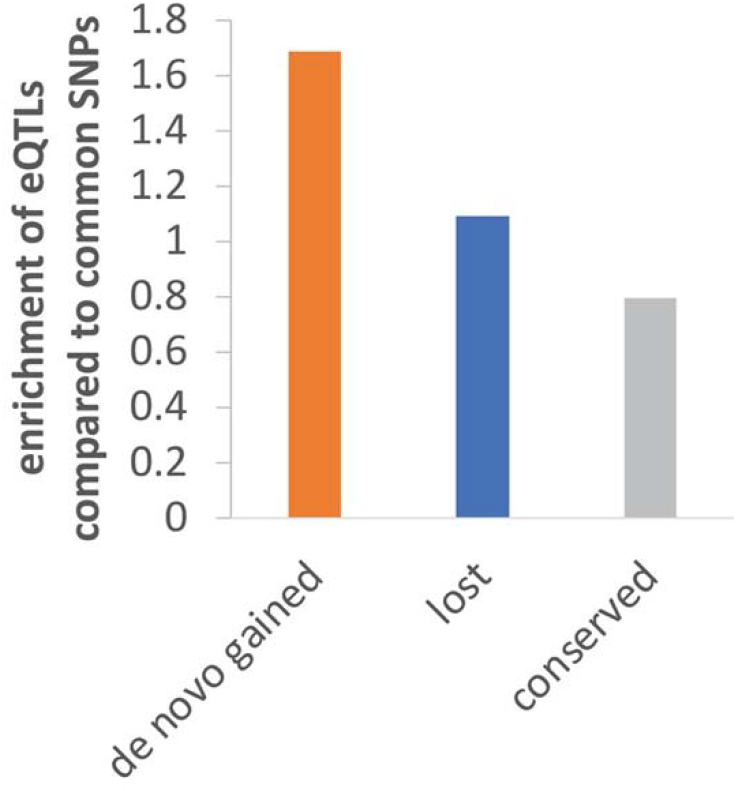
Enrichment of eQTLs compared to common SNPs in the three sets of enhancers. Specifically, the enrichment = fraction of eQTLs in enhancers/fraction of SNPs in enhancers.

**Figure S4.**
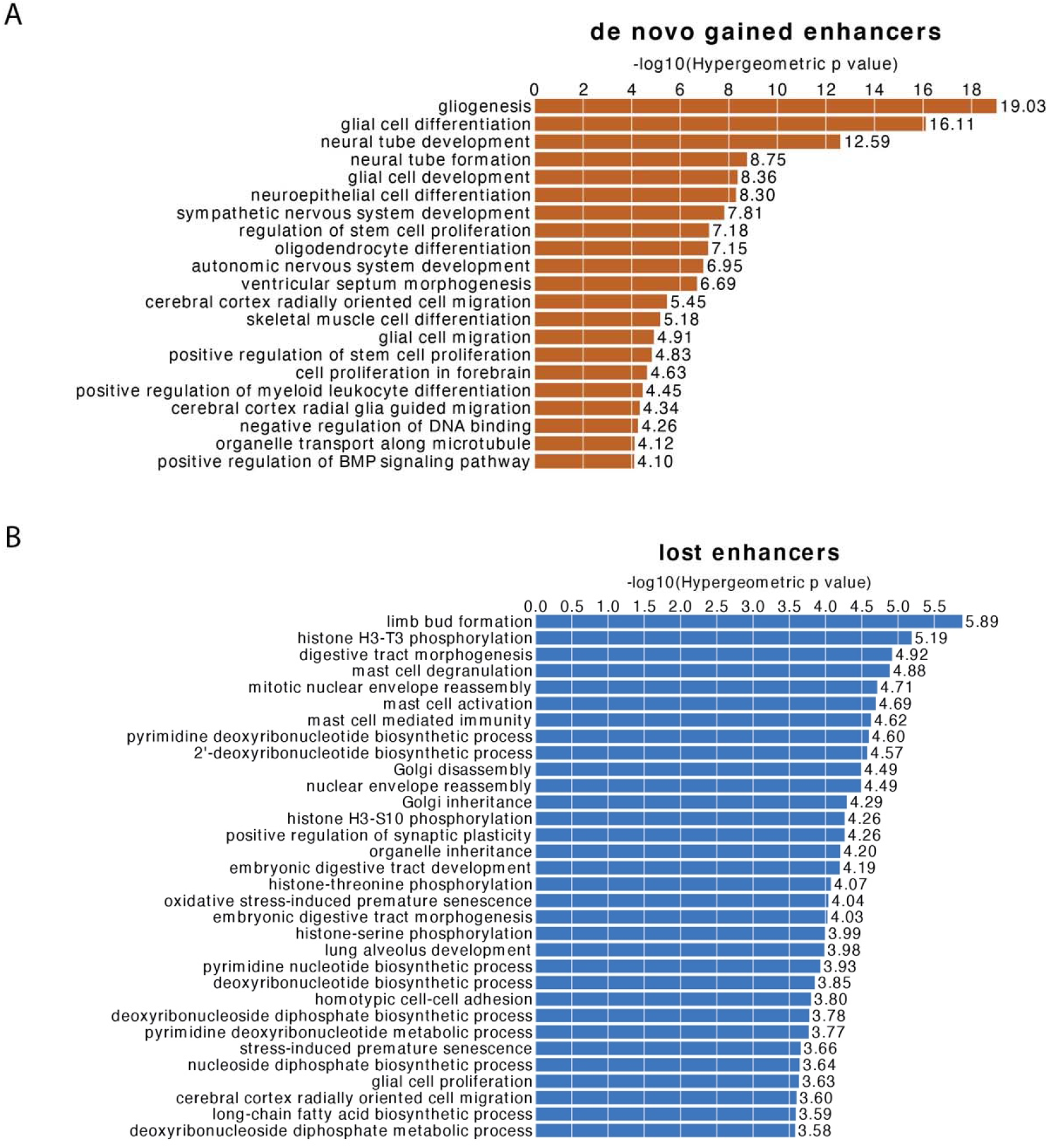
*De novo* gained enhancers are associated with essential CNS-related biological processes, using all fetal brain enhancers (de la Torre-Ubieta et al. 2018) as the background. (A) GO terms of *de novo* gained enhancers. (B) GO terms of lost enhancers. We apply GREAT with the single nearest gene association rule to do functional enrichment of genes near enhancers. The GO terms will be considered as enriched if it has at least 10 gene hits with FDR threshold set as 0.01.

**Figure S5.**
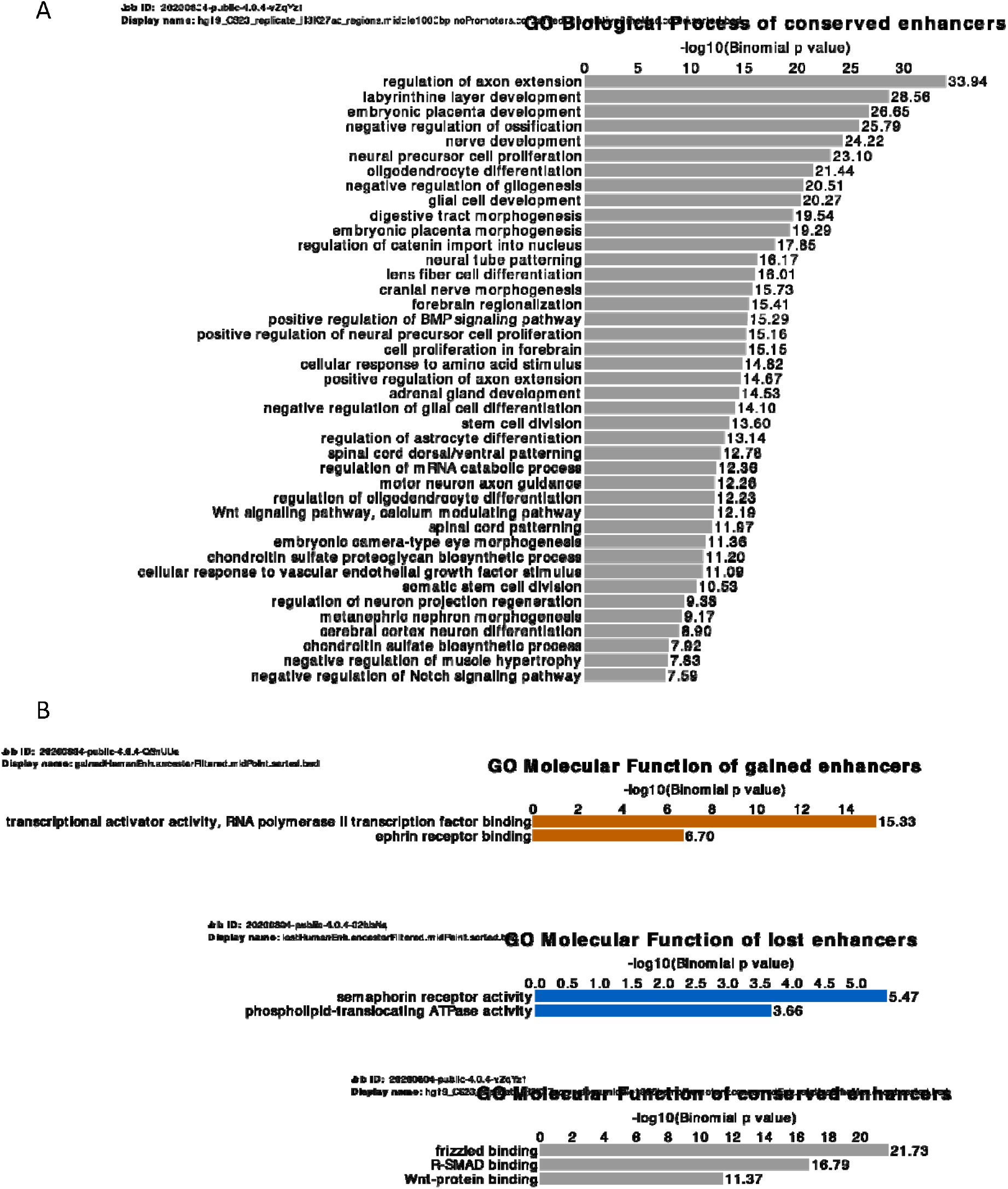
A) Enriched GO Biological Processes terms of conserved enhancers. B) Enriched GO Molecular Function terms of the three sets of enhancers.

**Figure S6.**
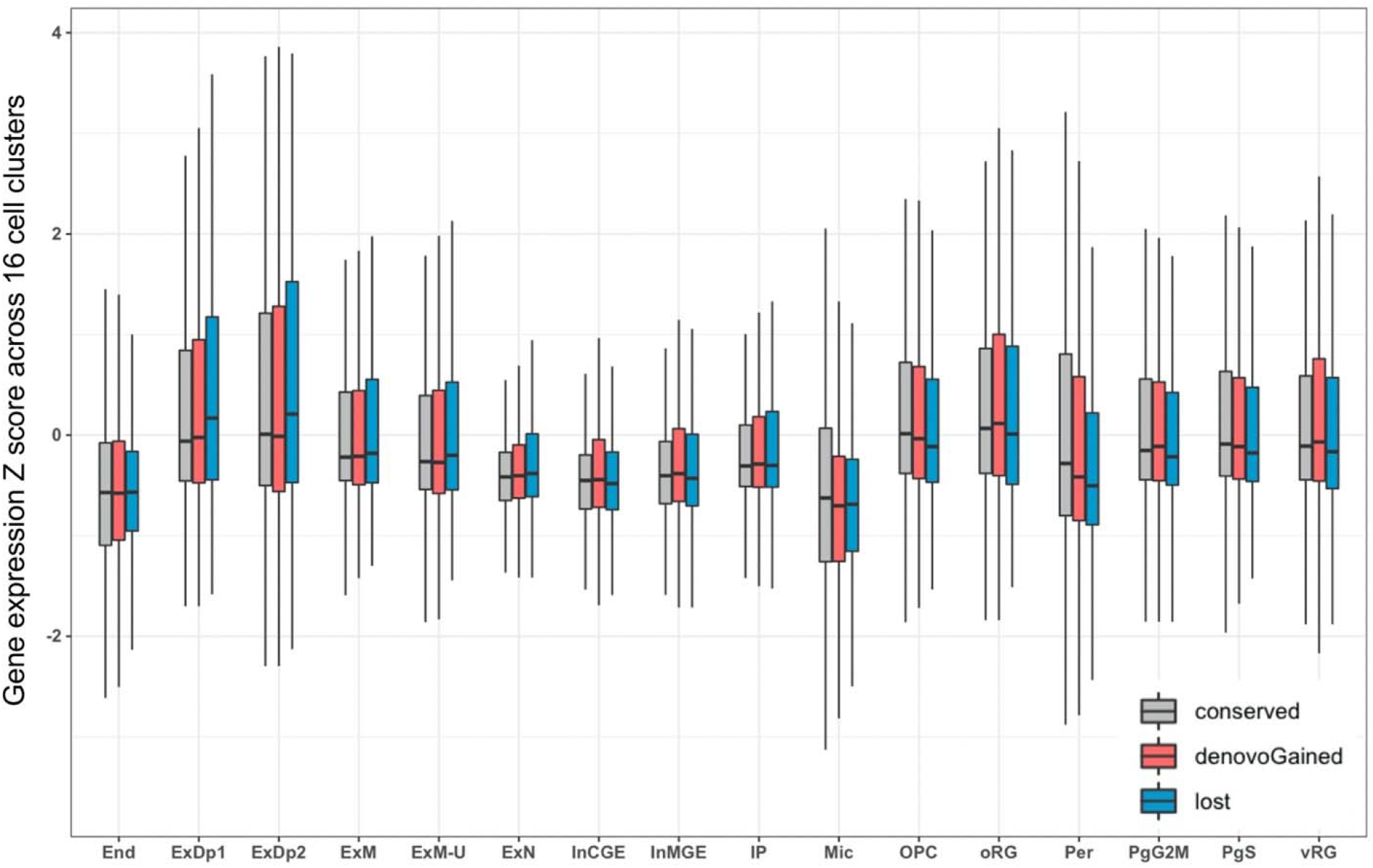
Z scores of expression of genes nearby the three sets of enhancers across 16 cell clusters. The lack of statistical significance may partly be due to the high variability/noise in single cell gene expression data, and also because only a subset of the genes near *de novo* gained enhancers are likely to drive cluster-specific expression as revealed in our fractional analysis (Figure 4C) but obscured in our analysis of z-scores for all genes.

**Figure S7.**
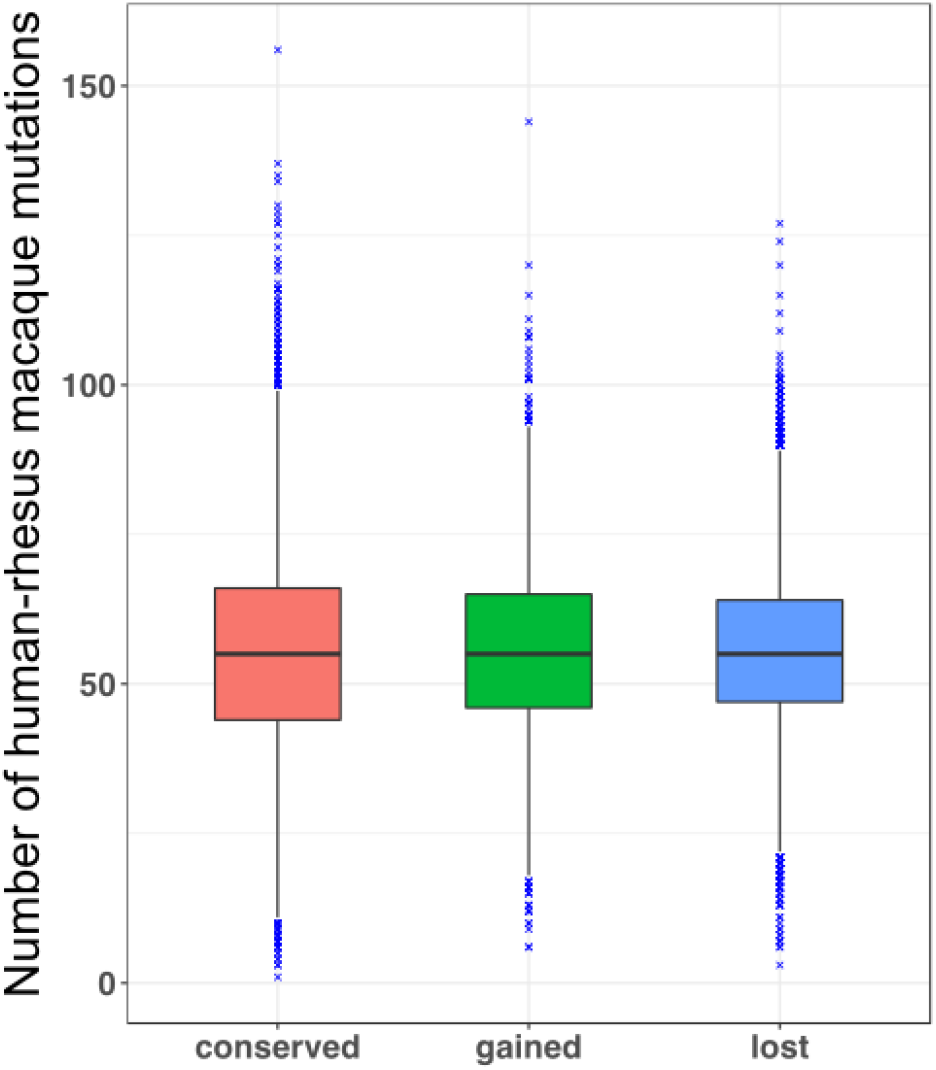
Number of human-macaque mutations within enhancers.

**Figure S8.**
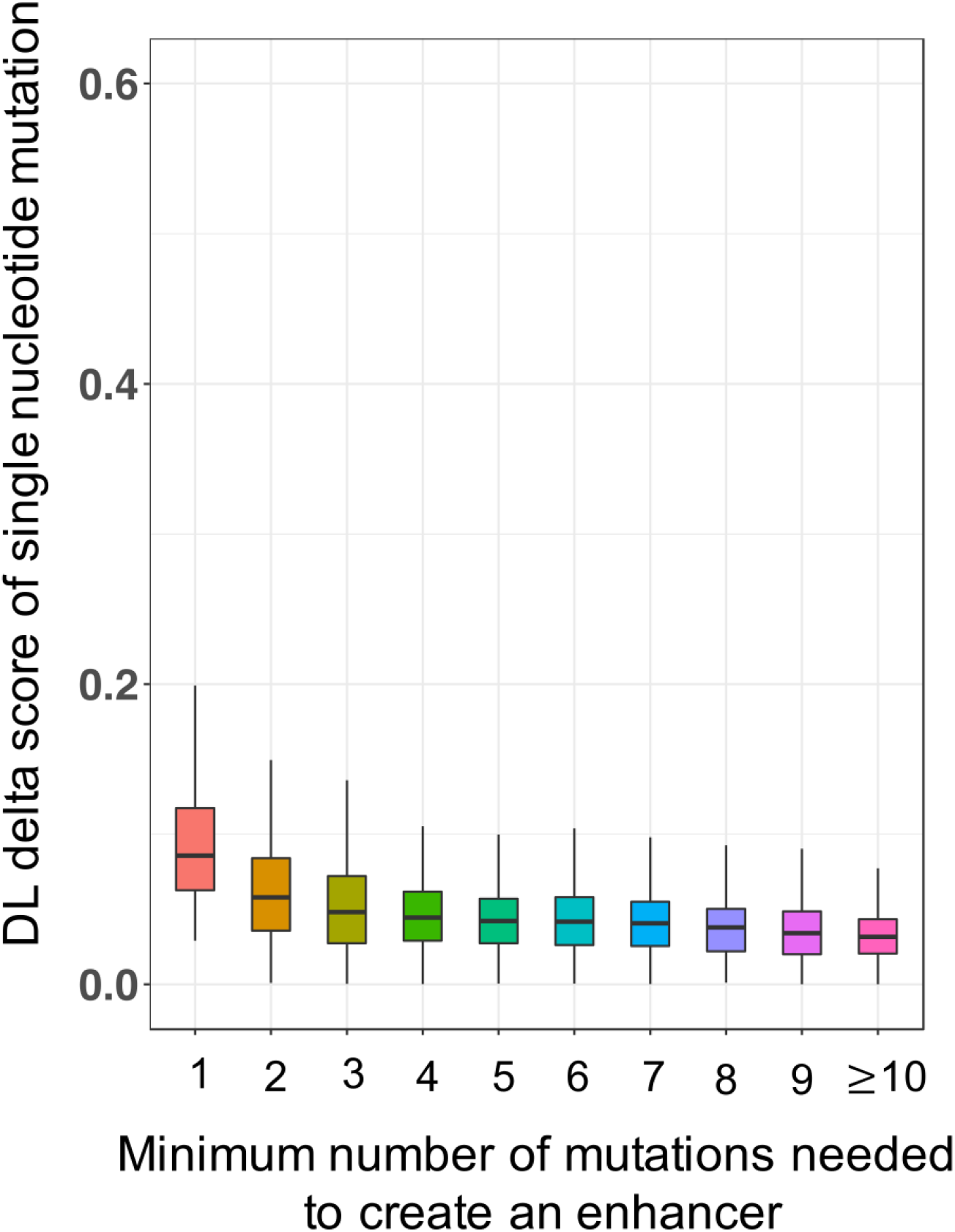
Distribution of delta score of the single nucleotide mutations that are minimally needed to create an enhancer.

**Figure S9.**
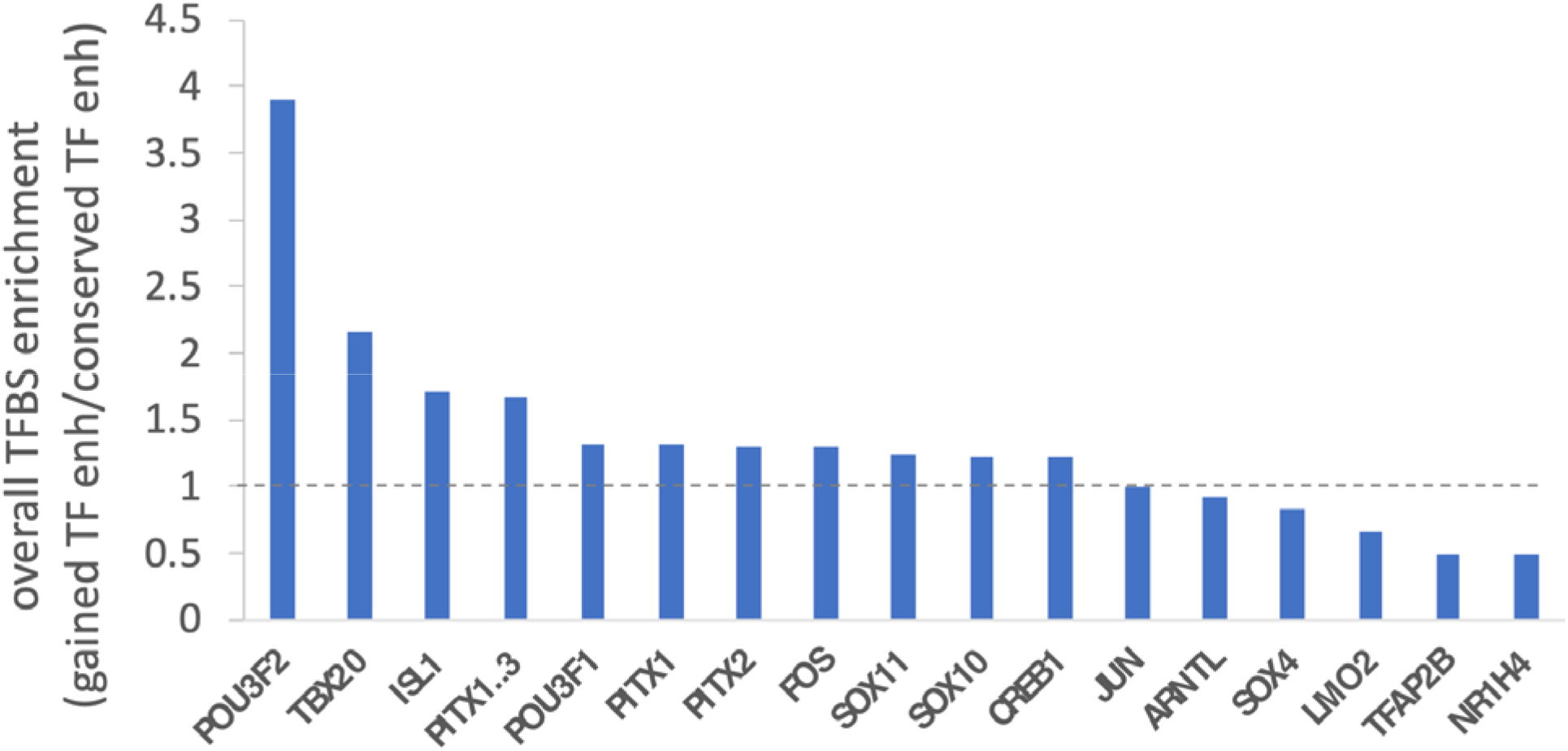
TFBS enrichment of gained enhancers associated with TFs, as compared to the conserved enhancers associated with TFs.

**Figure S10.**
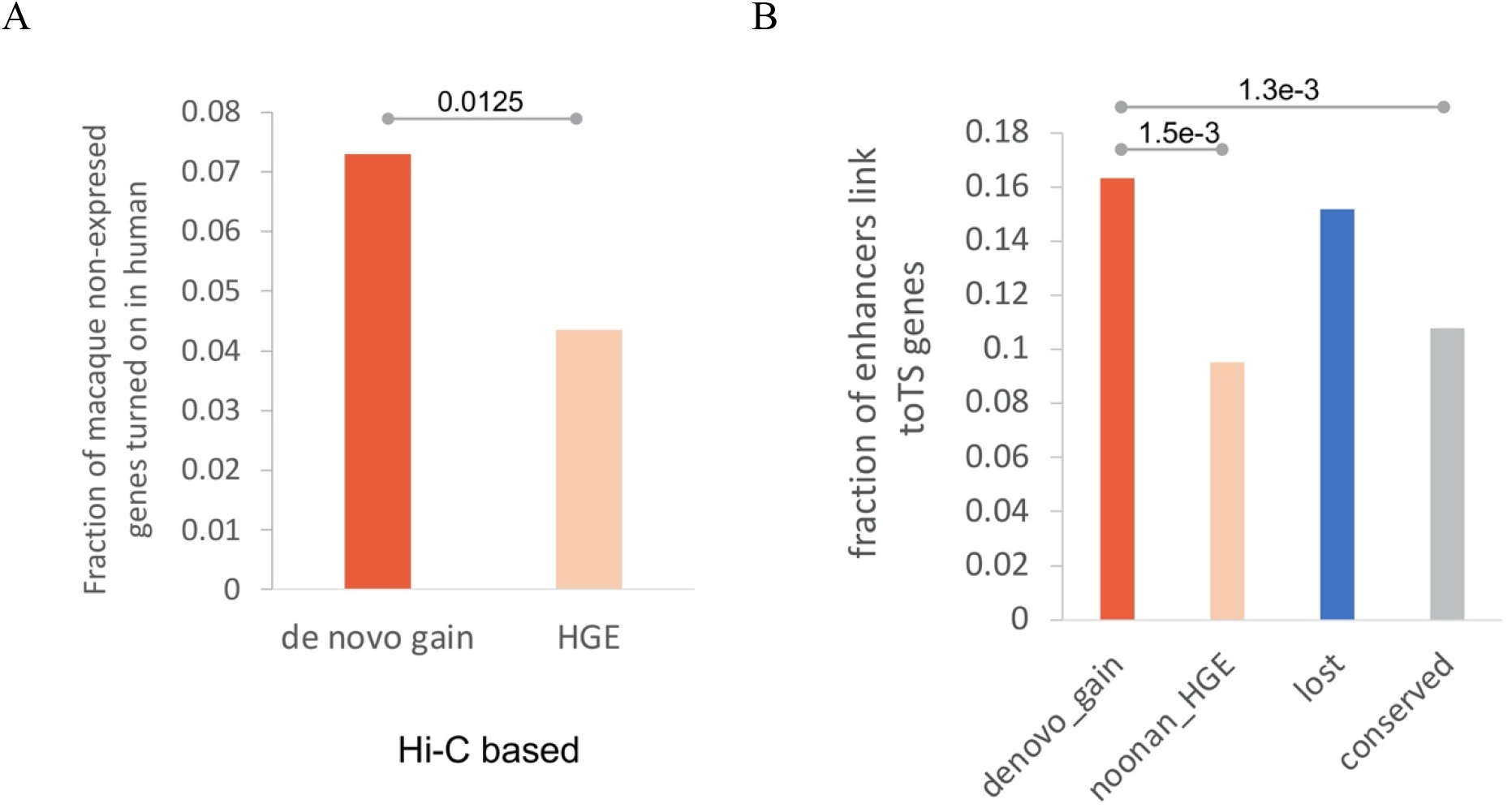
(A) Fraction of enhancers in contact with genes whose RPKM < 1 in macaque and > 1 in human. (B) Fraction of enhancers in 3D contact with the most tissue-specific genes.

**Figure S11.**
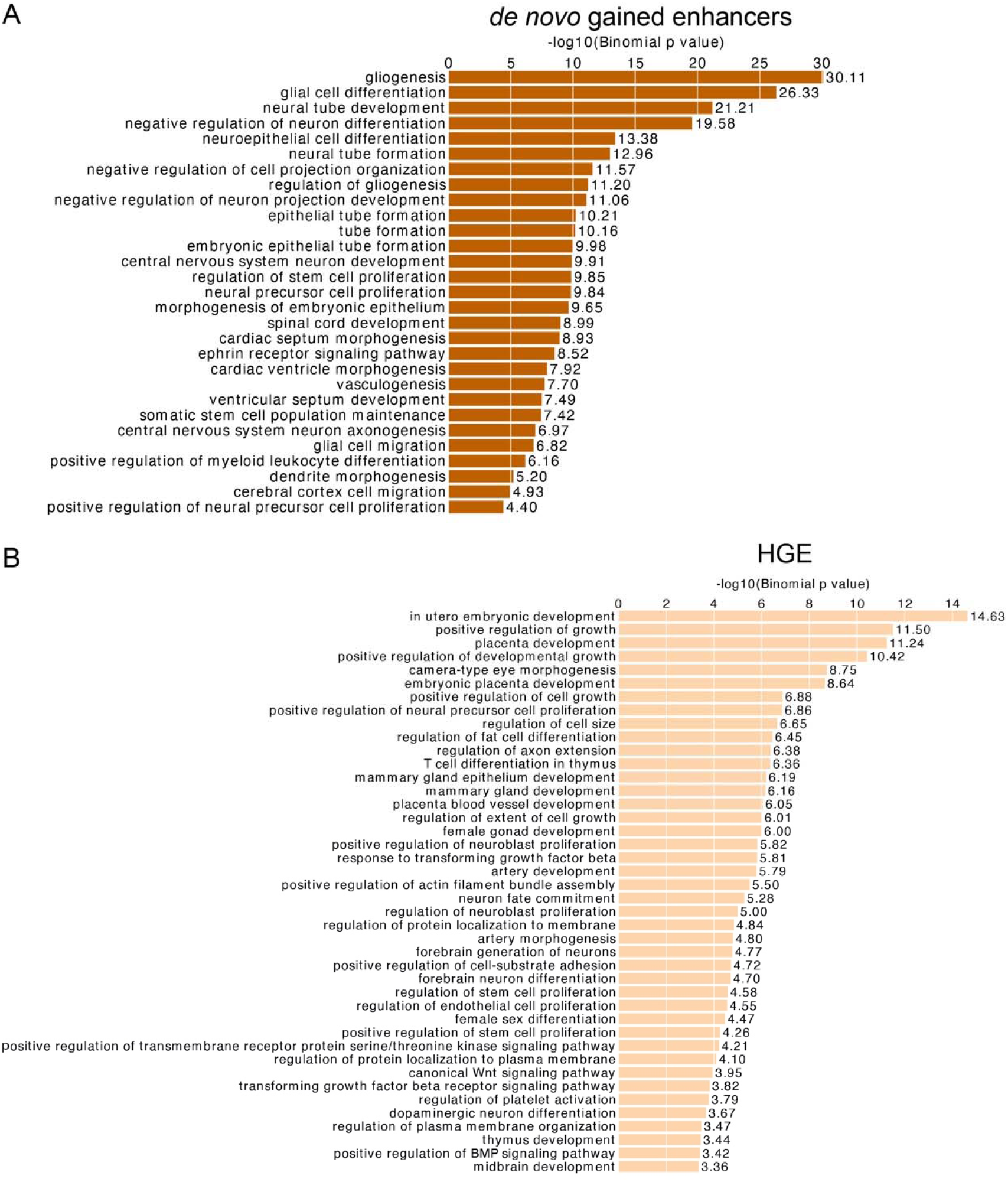
Enriched GO biological processes of *de novo* gained enhancers (A), and HGEs (B) using whole genome as the background.

**Figure S12.**
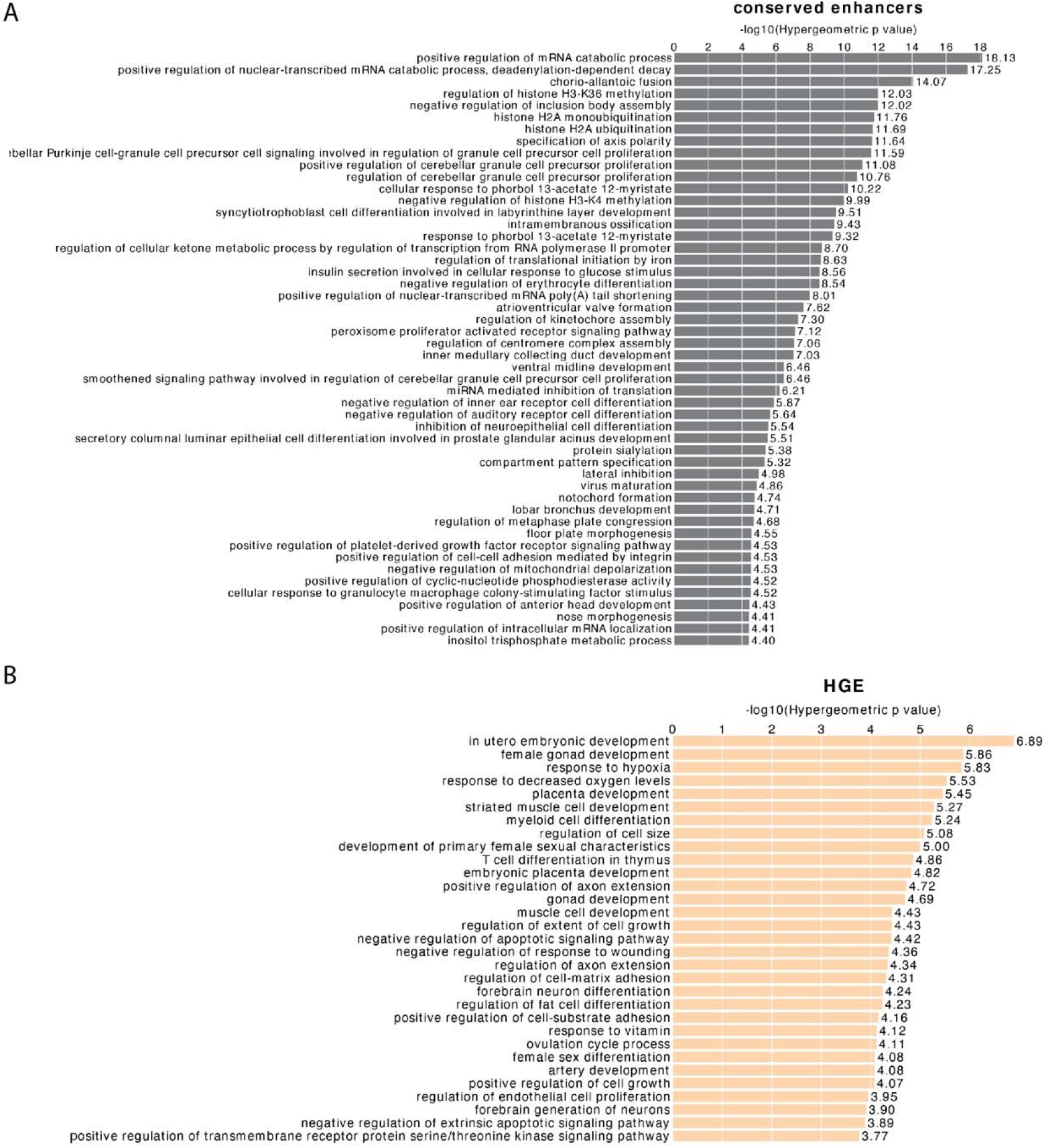
Enriched biological processes of a set of enhancers, using all fetal brain enhancers (de la Torre-Ubieta et al. 2018) as the background. (A) GO terms of conserved enhancers. (B) GO terms of HGEs. We apply GREAT with the single nearest gene association rule to do functional enrichment of genes near enhancers. The GO terms will be considered as enriched if it has at least 10 gene hits with FDR threshold set as 0.01.

**Figure S13.**
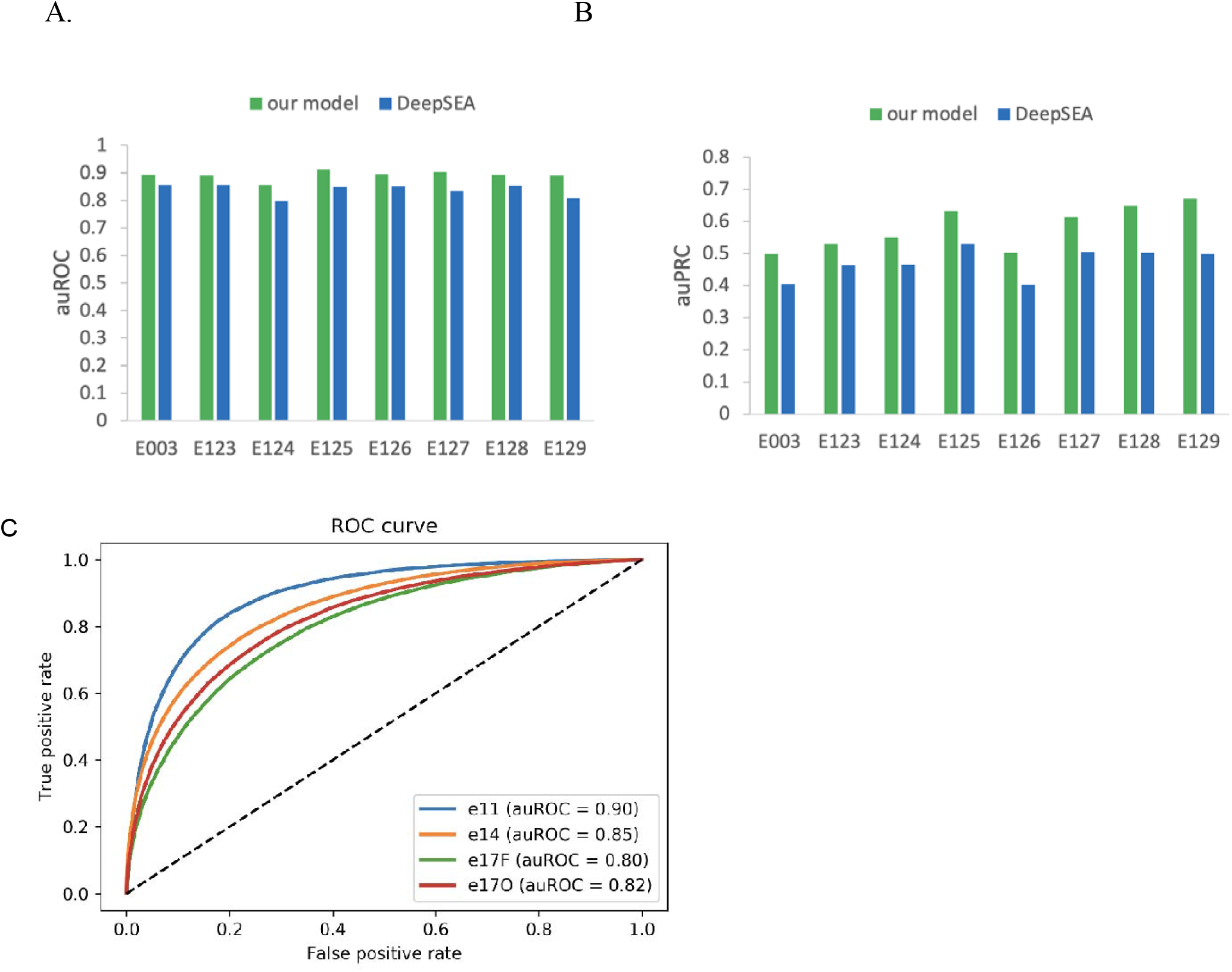
Performance of the DLM. (A) auROC and (B) auPRC of our model in predicting H3K27ac in 8 tissues which are tested by DeepSEA. (C). ROC curve of CS23 model tested on mouse embryonic neocortex enhancers corresponding to different stages of development (e11, e14, e17F, e17O). The E numbers on the x-axis are the tissue IDs defined by the Roadmap Epigenomic Project. E003: H1 Cell Line, E123: K562 Leukemia Cell Line, E124: Monocytes-CD14+ RO01746 Cell Line, E125: NH-A Astrocytes Cell Line, E126: NHDF-Ad Adult Dermal Fibroblast Primary Cells, E127: NHEK-Epidermal Keratinocyte Primary Cells, E128: NHLF Lung Fibroblast Primary Cells, E129: Osteoblast Primary Cells.

**Figure S14.**
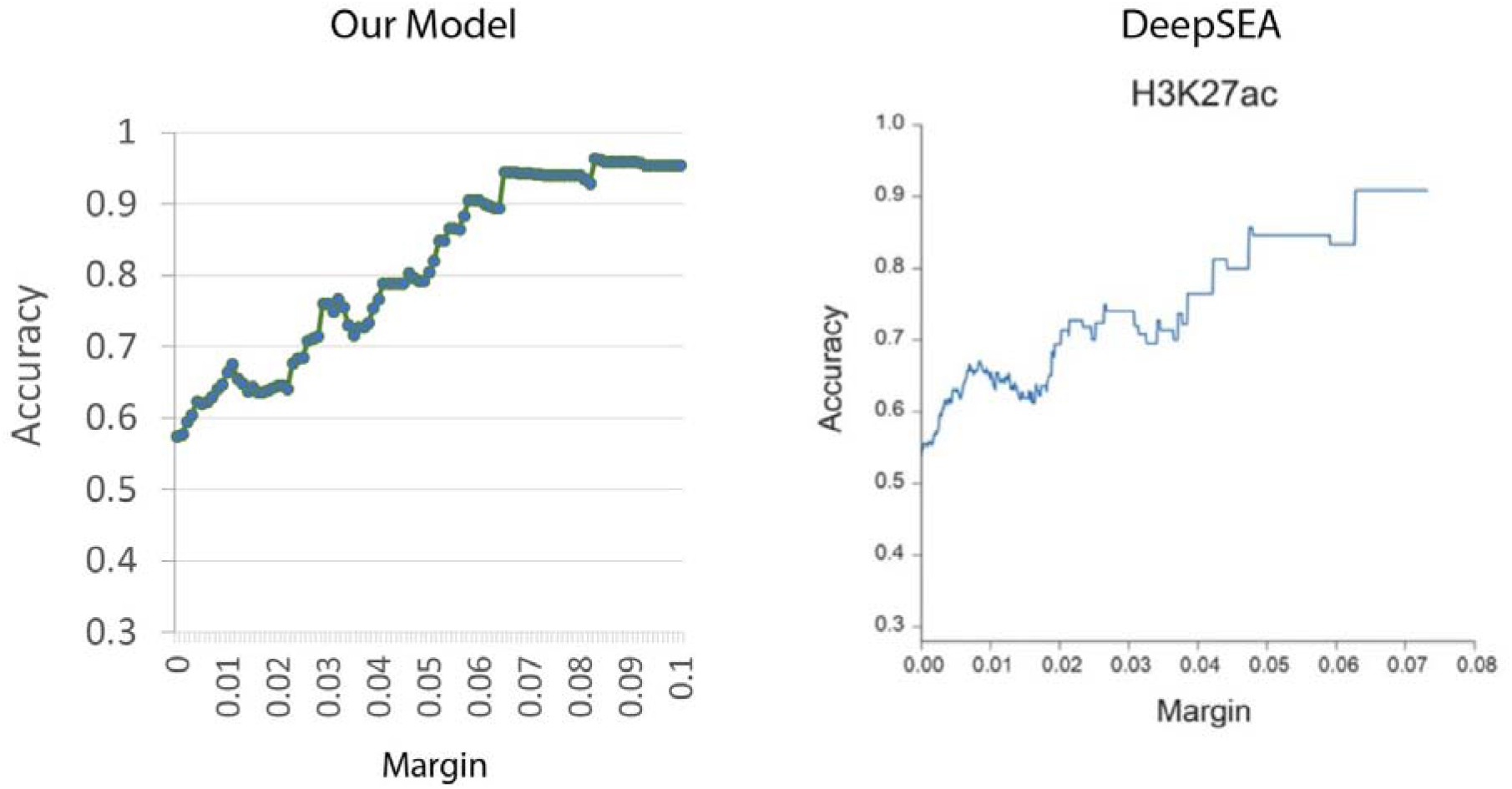
Deep learning histone mark classifiers provided accurate prediction of allele specific effects on histone marks H3K27ac (the allele with stronger histone mark signals). The predictions were evaluated with histone mark QTLs identified with FDR <0.1 in Yoruba lymphoblastoid cell lines (McVicker, G. et al. Science 342, 747-749 (2013)). Margin shown on the x axis is the threshold of predicted probability differences between the two alleles for classifying high-confidence predictions. Performance is measured by accuracy (y-axis) of predicting the allele with higher read counts based on DLM score difference above certain threshold (x-axis).

**Figure S15.**
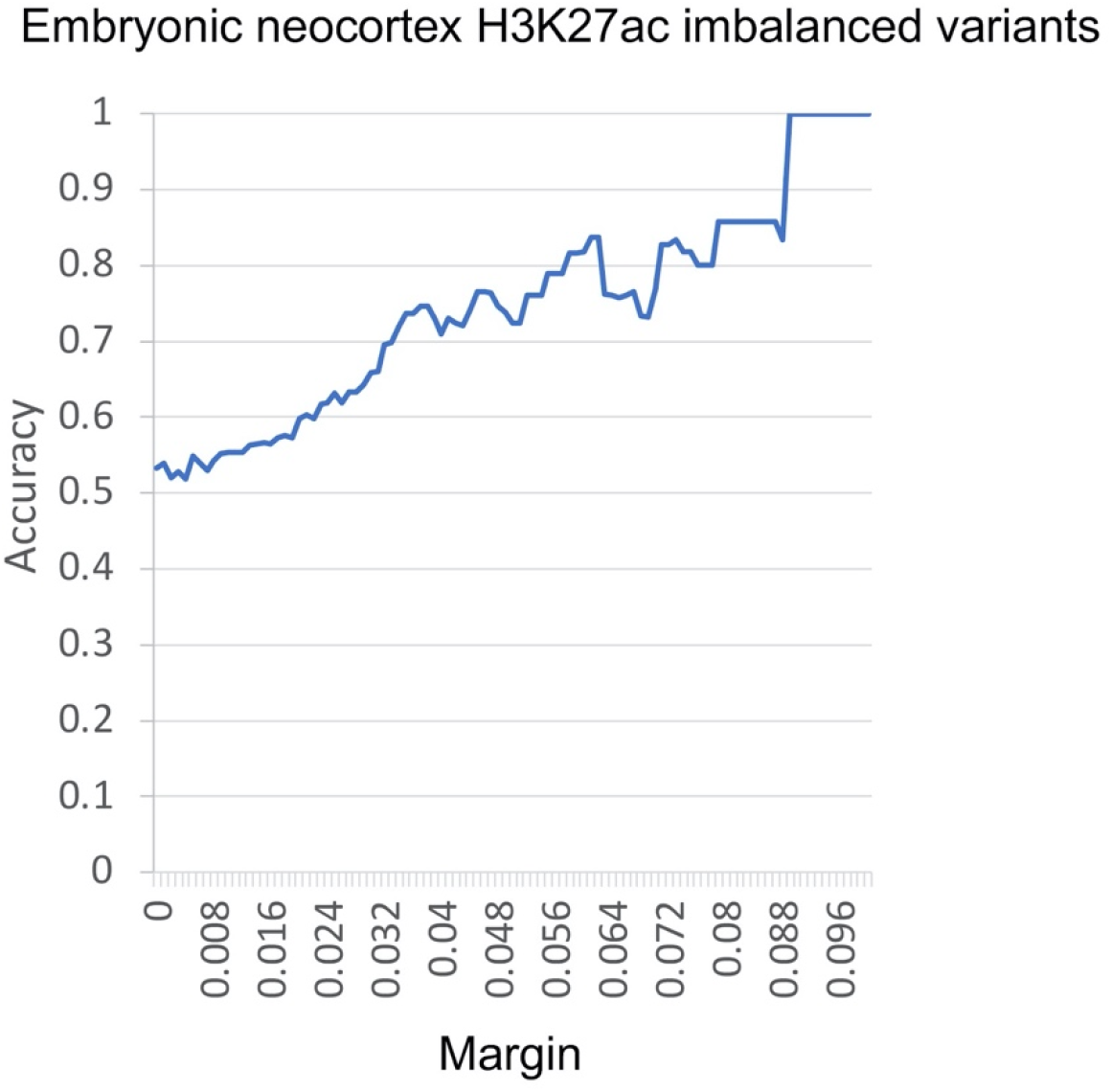
The DLM of CS23 H3K27ac accurately predict allelic imbalanced heterozygous variants within CS23 H3K27ac peaks.

**Figure S16.**
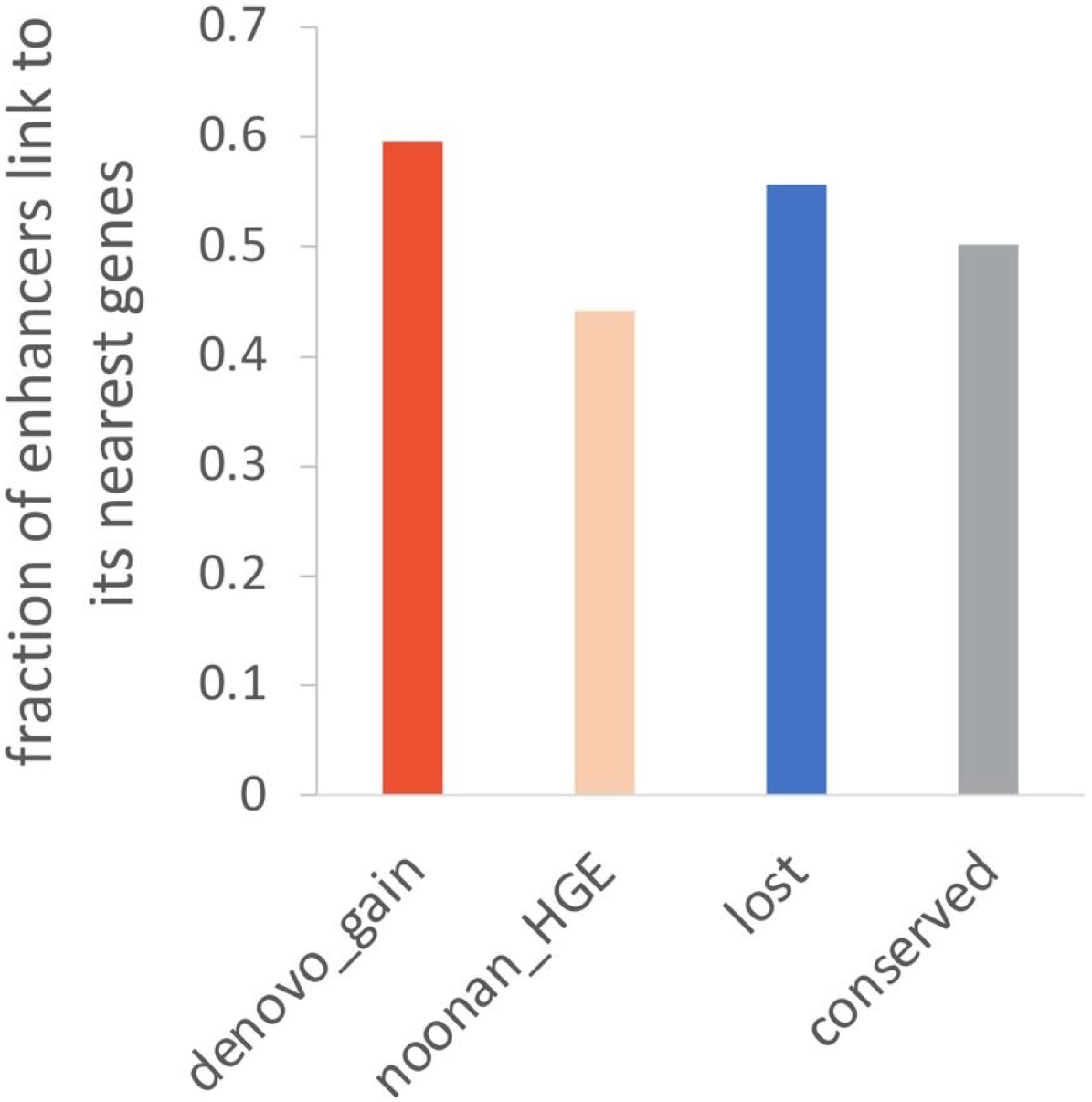
Fractions of enhancers that contact their nearest gene.

